# Structural modularity of the XIST ribonucleoprotein complex

**DOI:** 10.1101/837229

**Authors:** Zhipeng Lu, Jimmy K. Guo, Yuning Wei, Diana R. Dou, Brian Zarnegar, Qing Ma, Rui Li, Yang Zhao, Fan Liu, Hani Choudhry, Paul A. Khavari, Howard Y. Chang

## Abstract

Long noncoding RNAs are thought to regulate gene expression by organizing protein complexes through unclear mechanisms. XIST controls the inactivation of an entire X chromosome in female placental mammals. Here we develop and integrate several orthogonal structure-interaction methods to demonstrate that XIST RNA-protein complex folds into an evolutionarily conserved modular architecture. Chimeric RNAs and clustered protein binding in fRIP and eCLIP experiments align with long-range RNA secondary structure, revealing discrete XIST domains that interact with distinct sets of effector proteins. CRISPR-Cas9-mediated permutation of the Xist A-repeat location shows that A-repeat serves as a nucleation center for multiple Xist-associated proteins and m^6^A modification. Thus modular architecture plays an essential role, in addition to sequence motifs, in determining the specificity of RBP binding and m^6^A modification. Together, this work builds a comprehensive structure-function model for the XIST RNA-protein complex, and suggests a general strategy for mechanistic studies of large ribonucleoprotein assemblies.

## INTRODUCTION

Long noncoding RNAs (lncRNAs) play essential roles in many aspects of gene expression in development and disease (Wang and Chang, 2011). lncRNAs control X chromosome inactivation (XCI), genome imprinting, immune response, cell-cycle regulation, genome stability, lineage commitment and embryonic stem cell (ESC) pluripotency (Brannan et al., 1990; Brockdorff et al., 1991; Brown et al., 1991; Gomez et al., 2013; Guttman et al., 2011; Lee et al., 2016; Lu et al., 2017; Tichon et al., 2016). The list of functional lncRNAs is growing rapidly as more studies are conducted in a wide variety of biological systems. lncRNAs are distinguished from mRNAs in their processing, maturation, and ultimate mechanisms of action (Quinn and Chang, 2016; Ransohoff et al., 2018). Accumulating evidence has suggested that lncRNAs often serve as flexible scaffolds to recruit and coordinate multiple protein complexes to execute specific functions. For example, the yeast telomerase RNA recruits multiple proteins, and relocation of the protein binding motifs does not disrupt the function of the telomerase complex (Zappulla and Cech, 2004). The lncRNA HOTAIR recruits two distinct histone modification complexes, LSD1 and PRC2 to specify combinatorial patterns of histone modifications (Tsai et al., 2010).

Recently, several new experimental strategies have been developed and applied to lncRNAs to determine the structure and interactions that underlie their functions. In particular, Selective 2’ Hydroxyl Acylation analyzed by Primer Extension (SHAPE) and DMS-seq probe nucleotide accessibility, a proxy for RNA base pairing probability (Lu and Chang, 2016). These two approaches have been applied to several *in vitro* transcribed lncRNAs, such as XIST, HOTAIR, COOLAIR and Braveheart (Hawkes et al., 2016; Smola et al., 2016; Somarowthu et al., 2015; Xue et al., 2016). *In vivo* DMS-seq on the XIST RNA suggested functional local structure elements but did not reveal high level organization of the RNA (Fang et al., 2015). These studies have reported vaguely defined domains in these long transcripts, but it remains unclear whether these domains are relevant in physiological conditions. Computational modeling based on chemical probing is prone to errors especially for long transcripts (Eddy, 2004; Mathews et al., 2010).

Female placental mammals have two X chromosomes while males only have one. The difference in gene dosage relative to autosomes is compensated by a mechanism called X chromosome inactivation (XCI), which one of the two X chromosomes in females is randomly silenced. XIST (<u>X I</u>nactive <u>S</u>pecific <u>T</u>ranscript) is an essential lncRNA of ∼19kb that controls XCI by recruiting multiple proteins to deposit epigenetic modifications, remodel the X chromosome, and silence transcription in specific nuclear compartment (Brockdorff et al., 1991; Brown et al., 1991). Several recent studies used biochemical and genetic screens to find important new players in XIST functions (Chu et al., 2015; McHugh et al., 2015; Minajigi et al., 2015; Moindrot et al., 2015; Monfort et al., 2015). Xist associates with at least 81 proteins through direct RNA-protein and direct protein-protein interactions. If occurring all together, the XIST ribonucleoprotein complex is many times the size of the ribosome. A number of XIST-associated proteins mediate critical steps in XCI. For example, the RNA binding protein (RBP) SPEN binds the A-repeat of XIST and recruits the SMRT-HDAC3 complex to repress transcription (Lu et al., 2016; McHugh et al., 2015; Moindrot et al., 2015; Monfort et al., 2015). RBM15 and RBM15B recruit the WTAP-METTL3-METTLE14 (WMM) RNA methyltransferase complex to install m^6^A modifications, which are required for XIST function (Patil et al., 2016). LBR recruits the XIST-coated Xi to nuclear lamina for efficient silencing (Chen et al., 2016), HNRNPU family proteins and CIZ1 attach the XIST RNA to the inactive X chromosome (Hasegawa et al., 2010; Kolpa et al., 2016; Ridings-Figueroa et al., 2017; Sakaguchi et al., 2016; Sunwoo et al., 2017). A key question that remains is how are these numerous proteins and the structured XIST RNA assembled into a functional complex. Moreover, all of the above studies were done using mouse embryonic stem cells, and how the human XIST RNP may be organized given substantial sequence divergence is not known.

In our prior work, we used three orthogonal methods, PARIS, icSHAPE and structure conservation analysis to demonstrate a modular architecture of the XIST RNA. We found that the stochastic folding of the A repeat region serves as a multivalent platform for recruiting SPEN, a protein required for XCI. To determine the structural basis of the assembly of the multi-functional XIST RNP complex, we developed and integrated several structure-interaction analysis methods, including PARIS (psoralen crosslinking) (Lu et al., 2016), fRIP-seq (formaldehyde) (Hendrickson et al., 2016), eCLIP (UV crosslinking) (Van Nostrand et al., 2016), and PIRCh (purification of interacting RNA on chromatin, using glutaraldehyde crosslinking)(Fang et al., 2019). Together, we found that the entire XIST RNA-protein complex is folded in a modular manner. XIST-associated proteins cluster together in the 3D folded complex, instead of spreading along the linear sequence. The clustering of proteins on the XIST RNA structure predicts a modular organization of XIST functions. The folding of the RNA creates physical proximity that directs the m6A methylation complex and hence the modifications. Together, this structure-interaction analysis establishes a unifying model for XIST functions.

Using CRISPR-Cas9 genome editing, we reorganized the domain architecture of the XIST RNA by moving the A-repeat domain to other locations. This reorganization was followed by relocation of XIST-associated proteins, demonstrating a role of the architecture in separating the chromatin-binding and nuclear membrane binding regions of the XIST RNP, and a role of the domain architecture in guiding protein binding to the RNA, and the m^6^A modification of the RNA, which is required for XIST functions. Together, this study builds a comprehensive model for the XIST RNA-protein complex and establishes a paradigm for studying structural basis of lncRNA functions.

## RESULTS

### RNA chimeras in fRIP-seq reveal modular XIST RNP architecture

Using PARIS, icSHAPE and structure conservation, we have demonstrated that the XIST RNA is folded into modular and compact domains, each spanning hundreds to thousands of nucleotides (Lu et al., 2016). While a handful of the XIST binding RBPs have been mapped to distinct regions of the XIST RNA, the overall organization of this RNP complex remains unknown. We reasoned that, in addition to psoralen, other types of chemical crosslinkers, together with proximity ligation, could capture the compact higher order architecture of XIST, therefore providing additional lines of evidence for the XIST structure and RNA-protein interactions.

We searched published RNA-protein crosslinking studies and found RNA-RNA chimeras for XIST in a set of fRIP-seq (formaldehyde RNA immunoprecipitation sequencing) experiments targeting 24 RBP and chromatin-associated proteins in K562 cells, a female human myeloid cell line that undergoes proper X chromosome inactivation (Hendrickson et al., 2016; Lee et al., 2019; Minkovsky et al., 2015). Briefly, cells were lightly crosslinked with formaldehyde to fix RNA-protein interactions, sonicated to small fragments, and then RNA-protein complexes were immunoprecipitated (**Figure 1A**). It is important to recognize the caveat that the antibody used in fRIP may crossreact against additional RBPs, and the formaldehyde crosslinking may allow protein partners of the target RBP and their collective RNA cargos to be retrieved.

**Figure 1.**
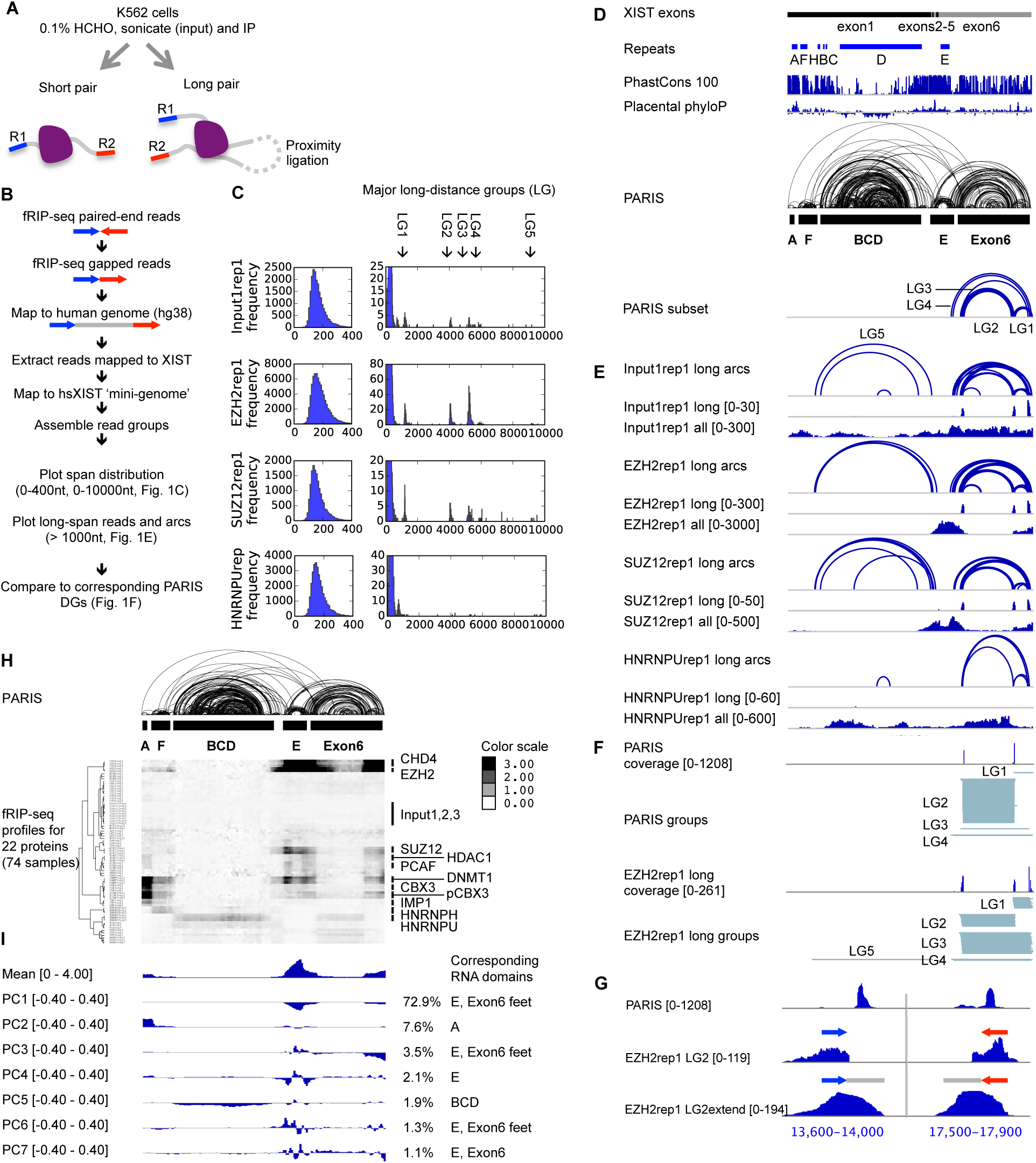
fRIP-seq confirms XIST RNA domains and reveals modular RNA-protein interactions. (A) Schematic diagram of the fRIP-seq experiment. The blue and red lines (R1 and R2, or read1 and read2), 31nt each, represent the paired-end sequenced tags. The gray lines represent the non-sequenced regions of the RNA fragments, each ∼200nt long. (B) Analysis strategy for the fRIP-seq data (see Supplementary Methods for details). Paired-end reads are rearranged and mapped to the genome using STAR, which reveals the non-sequenced fragment as a gap (gray line between the two sequenced tags R1 and R2). The mapped reads were remapped to the mature XIST RNA as a mini-genome to allow gap analysis and visualization. (C) Distribution of gaps or distances between the paired-end sequencing tags (R1 and R2). Most of the tag-pairs are from one RNA fragment, therefore within a short distance of each other (left side). A small fraction of the pairs are far away from each other, therefore most likely to be from two proximally ligated fragments (right side, same distribution, but highlighting the low-frequency long-distance pairs). Five major long pair groups (LGs) can be identified. The long distance distributions were plotted so that the y-axis is 1% that of the short distance distributions. The first biological replicate was shown for each protein and the average read numbers and standard deviations are calculated from all biological replicates (n=2 for EZH2 and n = 3 for the rest). (D) Annotation of the human XIST RNA. XIST exons and repeats, phylogenetic conservation (PhastCons 100 and Placental PhyloP from UCSC), and PARIS data in HEK293 cells are shown (Lu et al., 2016). Long distance groups that correspond to 4 fRIP-seq determined LGs (LG1-LG4) were extracted from PARIS data. (E) Long-distance arcs (tag pairs), coverage of long-distance tag-pairs, and coverage of all tags were shown for four examples (Input1, EZH2, SUZ12 and HNRNPU). These four samples correspond to the ones shown in panel (B). Y-axis scale is indicated in the square brackets. (F) Comparison of the overlapping duplex groups from PARIS and the long-distance groups from EZH2 fRIP-seq. (G) Comparison of the PARIS and EZH2 fRIP-seq long-distance group 2 (LG2), and LG2 extended to the average fRIP-seq fragment size (LG2extend [0-194]). Each side shows a 400nt window. (H) Unsupervised clustering of the XIST-Protein interaction profiles in 100nt windows. A total of 74 samples are clustered, excluding the samples STAG2 (non-specific, as determined by (Hendrickson et al., 2016)) and WDR5 (low read numbers). (I) PCA analysis of the profiles in 100nt windows, displaying the top 7 principal components, which explains more than 90% of all variation (top 4 components explain 86%). The percentages on the right represent variation explained by each component. See also Figure S1.

During the experiments, two adapters were ligated to the purified RNA fragments. In addition, the endogenous (in lysate) or the added RNA ligases (in purified RNA) can join two fragments that are crosslinked together, resulting in chimeras. Subsequent paired end sequencing captures the two ends of each chimera, and we developed a computational pipeline to identify such chimeras (**Figure 1B**). Short-distance pairs indicate single fragments, while long-distance pairs indicate two fragments that are proximally ligated in space. Distribution of distance between the paired-end sequencing tags (inclusive) is mostly between 100-400nts (**Figure 1C**, left side panels, see all distributions for all proteins in **Figure S1**). In addition, discrete clusters of long-distance reads are detected up to 10kb for most proteins, including the input control (**Figure 1C**, right side panels). Five major long-distance groups were identified in XIST. The discrete distribution suggests that the ligation reactions are highly specific for certain positions along the XIST RNA dictated by spatial proximity.

To determine the nature of these long-distance groups (LGs), we compared them to XIST structure model from PARIS experiments (Lu et al., 2016) (**Figure 1D**). The first 4 major LGs are all mapped to the exon6 domain as determined by PARIS, primarily among three anchor points, while LG5 is mapped to the large domain in exon1, encompassing repeats B, C and D (**Figure 1E**). LG1-4 overlap RNA duplexes from PARIS (**Figure 1F**); the sequencing tags extended to the approximate length of the RNA fragments, clearly overlaps the two arms of the PARIS duplex (**Figure 1G**). LG5 does not directly overlap duplexes in PARIS data but is consistent with the overall shape of the BCD domain. Together, these long-distance crosslinking data supports the XIST domain architecture as determined by PARIS, icSHAPE, and conservation analysis.

### fRIP-seq reveals spatial partition of XIST-associated proteins

To understand how XIST-associated proteins are assembled into the RNP, we normalized all 74 fRIP samples (including the input controls) against the average of input controls in 100nt windows (Supplemental Methods, the 25^th^ percentile set to 0.1 for each fRIP-seq profile), and clustered the enrichment profiles in an unsupervised manner (**Figure 1H**). Several patterns emerged from this analysis, including the selective enrichment in the A-repeat domain (DNMT1, CBX3 and phosphorylated CBX3 (pCBX3)), F domain (DNMT1, CBX3, pCBX3 and HNRNPH), BCD domain (HNRNPU, which is also enriched in the middle of the exon6 domain. This pattern was reproduced in the eCLIP analysis discussed below), E domain (CHD4, EZH2, SUZ12, DNMT1 and pCBX3), which is coupled to the end of the transcript. Together, these data show clear spatial separation of the XIST-associated proteins.

To automatically derive the domain definitions from the high dimension data, we applied principal component analysis (PCA, **Figure 1I**). The first 4 principal components (PCs) account for 86% of all variation, while the first 7 PCs account for >90% of all variation. The major domains are all detectable. For example, PC1 contains the coupled E-repeat and the “feet” of the Exon6 domain. PC2 primarily shows the different enrichment patterns in A and F domains. PC5 shows the differential enrichment in the largest BCD domain. Together these data not only confirm the XIST RNA structure but also reveal patterns of protein binding on the XIST RNA.

### Modular assembly of the XIST RNP based on eCLIP

To further understand the organization of the XIST RNP complex, we used published RNA-protein UV crosslinking data (eCLIP) to map the binding sites of XIST-associated proteins (Van Nostrand et al., 2016). Yeo and colleagues published a large set of RBP eCLIP data that maps the binding sites to nucleotide resolution. The original analysis revealed enrichment of only four proteins on the transcript level, including HNRNPM, HNRNPK, RBM15 and PTBP1, whereas many other XIST-associated proteins did not pass the stringent enrichment threshold. Out of the 81 proteins previously detected by Xist ChIRP in mouse cells (Chu et al., 2015), 27 of them are included in the 121 eCLIP dataset (**Table S1**).

Similar to the fRIP-seq, the existence of background renders the transcript-wide enrichment less obvious. To detect binding sites on XIST, we performed unsupervised clustering and PCA analysis on eCLIP data for 121 RBPs in K562 cells (**Figure 2**). Hierarchical clustering of the eCLIP profiles showed a pattern highly similar to the fRIP-seq data (**Figure 1H** and **2B**). The PCA analysis also revealed a pattern of RNA domains similar to the fRIP-seq, with slight differences in the intensity of the domains (**Figure 1I** and **Figure 2C**), although the top PCs explained less variation than the fRIP-seq data, likely due to the larger number of profiles (28.5% for eCLIP PC1 vs. 72.9% in fRIP-seq PC1). For example, PC1 corresponds to the proteins that preferentially bind the feet of the Exon6 domain, the same as in fRIP-seq, while PC2 and PC3 corresponds to enrichment primarily on the A and F domains. The middle regions of the BCD and Exon6 domains are depleted of most protein binding. Together, these data not only support the domain architecture of the XIST RNA, but also revealed clustered binding of the proteins (**Figure 2D**). We named each of the major structural domains of XIST after the primary sequence repeat that is present in the corresponding domain (“domains A, F, BCD, E”), except the very 3’ “Exon6 domain” which contains no sequence repeats. Salient features for three of the domains are presented below, and a subset of the associated proteins are discussed. Similar analysis of two other lncRNAs, MALAT1 and NEAT1 also revealed modular domains, although the domain definitions are not as clear as XIST (**Figure S2A-F**).

**Figure 2.**
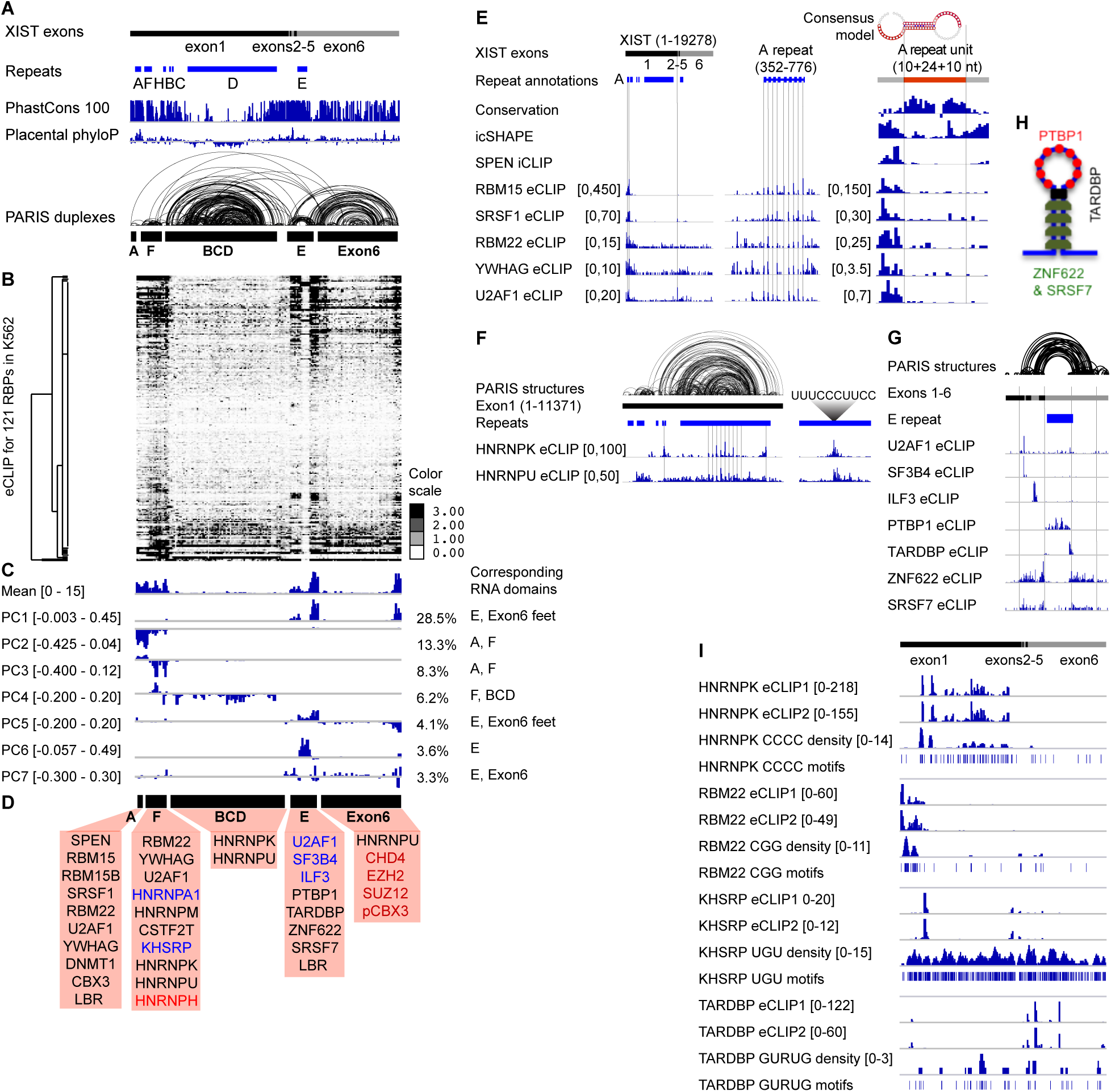
eCLIP analysis of XIST-associated proteins reveal modular domains. **(A)** Annotation of the human XIST RNA. XIST exons and repeats, phylogenetic conservation (PhastCons 100 and Placental PhyloP from UCSC), and PARIS data in HEK293 cells are shown, same as Figure 1D. **(B)** Unsupervised clustering of XIST-protein interaction profiles in 100nt windows for all 242 samples of the 121 proteins, two biological replicates each. **(C)** PCA analysis of all eCLIP data in 100nt windows. The mean and first 7 principal components are displayed together with percentage of variation explained by each component on the right. **(D)** List of proteins associated with each domain. Black and blue letters: enriched proteins based on eCLIP. Black: broad and clustered binding. Blue letters: focal binding. Red letters: enriched proteins based on fRIP-seq. **(E)** RBP interaction data for the A domain. The left panel shows enrichment of 6 RBPs along the entire XIST RNA based on eCLIP. The three vertical lines highlight the A-repeat region and the SRSF1 and U2AF1 peaks on exon 2. The middle panel shows zoom-in to the A-repeat domain and the vertical lines indicate the crosslinking positions of the six proteins on the repeat units. The right panel shows the average signal on a single 24nt repeat with 10nt flanking spacer sequences on each side, and the vertical lines mark the start and end of the 24nt repeat sequence. Conservation, icSHAPE and SPEN iCLIP data were from (Lu et al., 2016). One replicate was shown for the eCLIP data of each protein. **(F)** RBP interaction data for the BCD domain. In the left panel, human 293T cell PARIS data, repeat annotations and eCLIP data are shown for the entire exon 1 (Lu et al., 2016). Vertical lines mark the boundaries of the BCD domain and the internal repetitive sequences in the D-repeat. The right panel shows the average of all D repeats (290nt per unit) and the consensus HNRNP binding site. One replicate was shown for the eCLIP data of each protein. **(G)** RBP interaction data for the E domain. Parts of exons 1 and 6, and the entire exons 2-5 are shown, together with human 293T PARIS data (Lu et al., 2016). The vertical lines mark the boundaries of the “stem” and “loop” regions of this giant stemloop. One replicate was shown for the eCLIP data of each protein. **(H)** Schematic RBP interaction model for the E domain. Only the four proteins that mark the “stem” and “loop” are shown. **(I)** Comparison of protein binding sites on XIST based on eCLIP and the sequence motifs. eCLIP data are normalized against size-matched input in 100nt windows. The density profile was calculated based on sequence motifs in 300nt windows and 50nt steps. See also Figure S2.

#### XIST A domain

The A-repeat region is essential for the silencing activity of XIST (Wutz et al., 2002). We have previously found that SPEN specifically binds the A-repeat (Lu et al., 2016). In addition, eCLIP and fRIP nominates multiple additional proteins that bind the A-repeat (**Figure 2D**, and **2E**, left panel, see **Figure S2G** for RBP enrichment distribution). SPEN and RBM15, RBM15B are in the same family, each having similar N-terminal RRM domains and a C-terminal SPOC domain. SRSF1, RBM22 and U2AF1 are all directly involved in splicing. Both SRSF1 and U2AF1 showed clear interaction with Exon2, consistent with their possible roles in splicing during XIST biogenesis (but no clear interaction with XIST introns, data not shown). The folding up of F domain and BCD domain brings the A-repeat region close to the internal exons 2-5, suggesting a role of the high level architectures in regulation of splicing. Panning and colleagues identified SRSF1 as an essential A-repeat-associated factor for efficient XIST splicing (Royce-Tolland et al., 2010). Our analysis thus provides a potential explanation for how the distant binding at the A-repeat could affect the splicing of the internal exons. Interestingly, all 6 proteins are crosslinked to the A repeat in a periodic fashion, primarily in the single-stranded spacer regions at the junction of double-stranded duplexes formed by hybridization of the repeat subunits (**Figure 2E**, middle panel, vertical lines). Averaging the repeats showed that the primary crosslinking sites are in single-stranded the spacer region, with slight differences among the 6 proteins (**Figure 2E**, right panel). The high affinity of multiple proteins to the A-repeat domain suggests that this domain acts as a nucleation center for XIST RNP assembly.

#### XIST F domain

Multiple proteins are enriched in the F domain, including the splicing factors U2AF1 and RBM22, and several HNRNP proteins. A function for the F domain in X inactivation has not been described. The close proximity to the A and BCD domains may explain the association with similar protein factors.

#### XIST BCD domain

The middle of the two large domains, BCD and Exon6 are generally depleted of protein binding except HNRNPK and HNRNPU based on the fRIP-seq and eCLIP (**Figure 1H-I** and **Figure 2B-C**, note the lower signal in both “Mean” tracks). HNRNPK binds primarily to the BCD domain to multiple clusters, while HNRNPU binds F, BCD and middle of Exon6 domains, also in clusters (**Figure 2F**, left panel). The human XIST RNA contains 14 units of the D repeat, which explains the periodical binding patterns for HNRNPK and HNRNPU. Averaging eCLIP signal for these two proteins revealed a clear pattern and the pyrimidine-rich consensus sequence (**Figure 2F**, right panel). HNRNPU binding is more broad than HNRNPK, especially in the repBCD domain, where the HNRNPU is widely dispersed while HNRNPK is more concentrated on a few peaks. Genetic deletion of mouse Xist BC regions abrogates hnRNPK binding and the associated PRC1 complex, validating this finding (Bousard et al., 2019).

#### Xist E domain

The E domain contains the E repeat, a highly degenerate pyrimidine rich region, and its surrounding flanking sequences (∼600nt each side). Seven proteins with strong binding sites on the E domain are discovered based on the eCLIP data (**Figure 2G**). Two splicing factors, U2AF1 and SF3B4 bind a focal point in Exon2, while the other 5 proteins bind broad regions. ILF3, a double strand RNA binding protein (dsRBP) bind the evolutionarily conserved exon4, which is required for high level XIST expression (**Figure S3**) (Caparros et al., 2002). The focal binding of ILF3 to exon4 is highly significant, but was masked by the background when performing whole-transcript enrichment analysis, and therefore was not considered as enriched in the original analysis (Van Nostrand et al., 2016). PTBP1 binds the E repeat region in the E domain, consistent with its preference for pyrimidine rich sequences (**Figure 2G, H**). TARDBP binds the junctions between the unstructured E repeat and the two stems, or the “neck” of the giant E domain stemloop (**Figure 2G, H**). ZNF622 and SRSF7 bind the stem regions, contrary to PTBP1. Together four proteins, PTBP1, TARDBP, ANF622 and SRSF7 show clear spatial partition on the E domain (**Figure 2H**).

#### XIST Exon6 domain

Similar to the large BCD domain, the center of the Exon6 domain is generally depleted of protein binding. The most significantly associated protein is HNRNPU, which was detected in both fRIP and eCLIP. Interestingly, the two sides, or feet, of the Exon6 domain associated with multiple proteins. This bimodal placement is consistent with the overall structure of the Exon6 domain, which brings the two ends to physical proximity.

#### Sequence and structure specificity of RBP binding

The clustered binding sites that correlate with high-level structural features suggest that RNA structures contribute to binding specificity. To understand how RBP specificity is determined, we analyzed the correlation between sequence motifs and actual binding sites from eCLIP experiments (Dominguez et al., 2018; Van Nostrand et al., 2017). For HNRNPK and RBM22, the eCLIP read density closely follows motif density, while for KHSRP and TARDBP, there is very little correlation, suggesting that additional factors contribute to the RBP specificity (**Figure 2I**).

### Conservation of the XIST domain architecture in mammals including mouse

Using PARIS, icSHAPE, and PARIS-guided multiple sequence alignments, we previously found that the XIST structure domains are highly conserved in evolution, despite limited sequence conservation (Lu et al., 2016). To further confirm the structure conservation in mouse, we performed PARIS in mouse ES cell line HATX3 (*Xist^TX/TX^ Rosa26^nlsrtTA/nlsrtTA^*), expressing Xist from the endogenous locus under a tetracycline-inducible promoter (Monfort et al., 2015) (**Figure 3A**). Despite the lower sequencing coverage, we detected 108 duplex groups after lifting to the human XIST coordinates, and found that all the previously discovered major domains are present in mouse Xist. To directly compare XIST structures between human and mouse, we lifted mouse PARIS data to human XIST coordinates (**Figure 3B**). About 42% of mouse Xist DGs overlap with human XIST DGs from HEK293 cells (Lu et al., 2016) (1000 times shuffling of all duplexes, p value < 0.001), suggesting that the overall Xist structure is conserved between human and mouse, despite the major differences in the size of repeats (Elisaphenko et al., 2008). Therefore, we conclude that the overall architecture of XIST is conserved in evolution. One of the most highly conserved long-range duplexes, connecting the start and end of the BCD domain, is very stable, with 37 closely stacking base pairs (minimum free energy = −31.70 kcal/mol, **Figure 3C**, see also **Figure 6D** from (Lu et al., 2016)).

**Figure 3.**
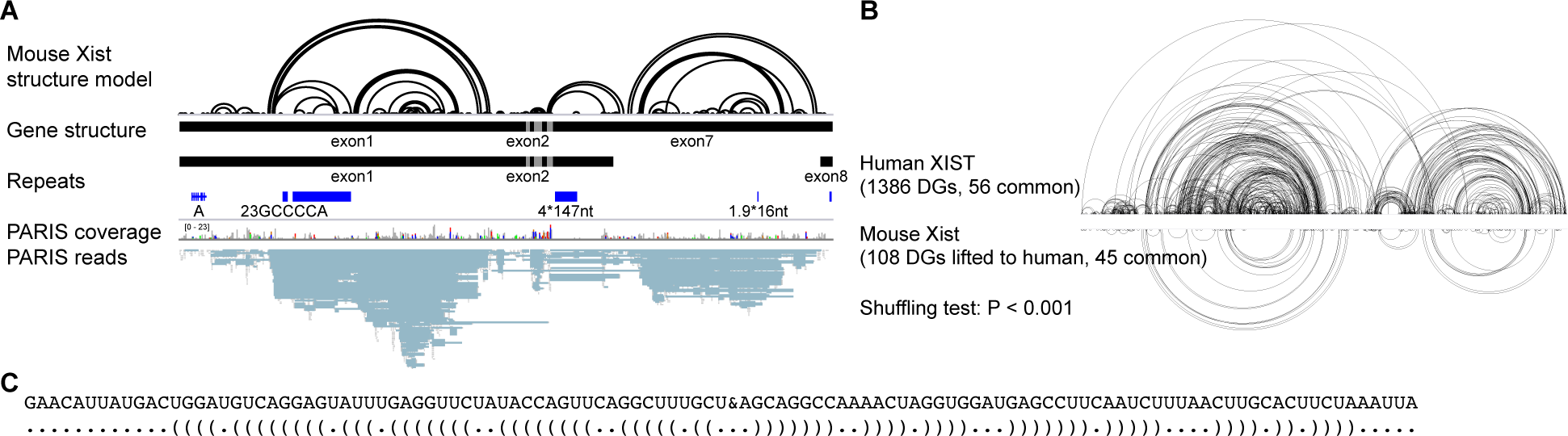
Conservation of the XIST architecture in human and mouse. (A) PARIS analysis of mouse Xist RNA structure in HATX mES cells. The structure model is the duplexes detected by PARIS. The gene structure tracks are the two mouse mature Xist isoforms. Mouse Xist repeats were detected using the Tandem Repeat Finder (Benson, 1999). (B) Comparison of human and mouse Xist PARIS-determined structures. The mouse Xist DGs are lifted to human coordinates for comparison with the human XIST PARIS data from 293T cells. (C) A highly conserved long range duplex structure as detected by PARIS in human and mouse (likely the ‘boundary element’)

### The domain architecture of XIST RNP determines m^6^A modification specificity

XIST RNA contains a large number of m^6^A modifications in both human and mouse cells (Linder et al., 2015; Patil et al., 2016). m^6^A sites closely follows RBM15/RBM15B occupancy, which recruit the METTL3/14 methyltransferase complex, suggesting that these two adapter proteins guide the modification. Patil et al. proposed a model where the location of RBM15/15B proteins determines m^6^A modification sites. Close examination of CLIP data showed that the RBM15/RBM15B binding sites are primarily clustered in the A-repeat domain and the other sites are much weaker, raising the question of how m^6^A is placed at distal locations on XIST RNA over ten thousand bases away. Instead, we noticed that all m^6^A modification sites, as well as the RBM15/RBM15B binding sites, are in close spatial proximity when the XIST RNA is folded (**Figure 4A**). The consensus m^6^A motif DRACH is nearly uniformly distributed along the XIST RNA transcript in both human and mouse, with the exception of the pyrimidine-rich E repeat, and at a density of one motif every ∼54nt (**Figure 4A**). The discrepancy between m^6^A motifs and modification sites suggests that the folding of the XIST RNA into compact modular domains contribute to the specificity of m^6^A modifications. Here we propose two hypotheses to explain the pattern of m^6^A modifications. First, the compact folding of the large BCD and Exon6 domains exclude the m^6^A methylase complex. Second, the folding of the XIST RNA creates local proximity among the modification sites with the A-repeat domain. These two possibilities are not mutually exclusive. Altering the overall structure of the XIST RNA is challenging because all the base pairing interactions contribute to the whole transcript structure; disruption of a small number of base pairs is unlikely to cause global changes.

**Figure 4.**
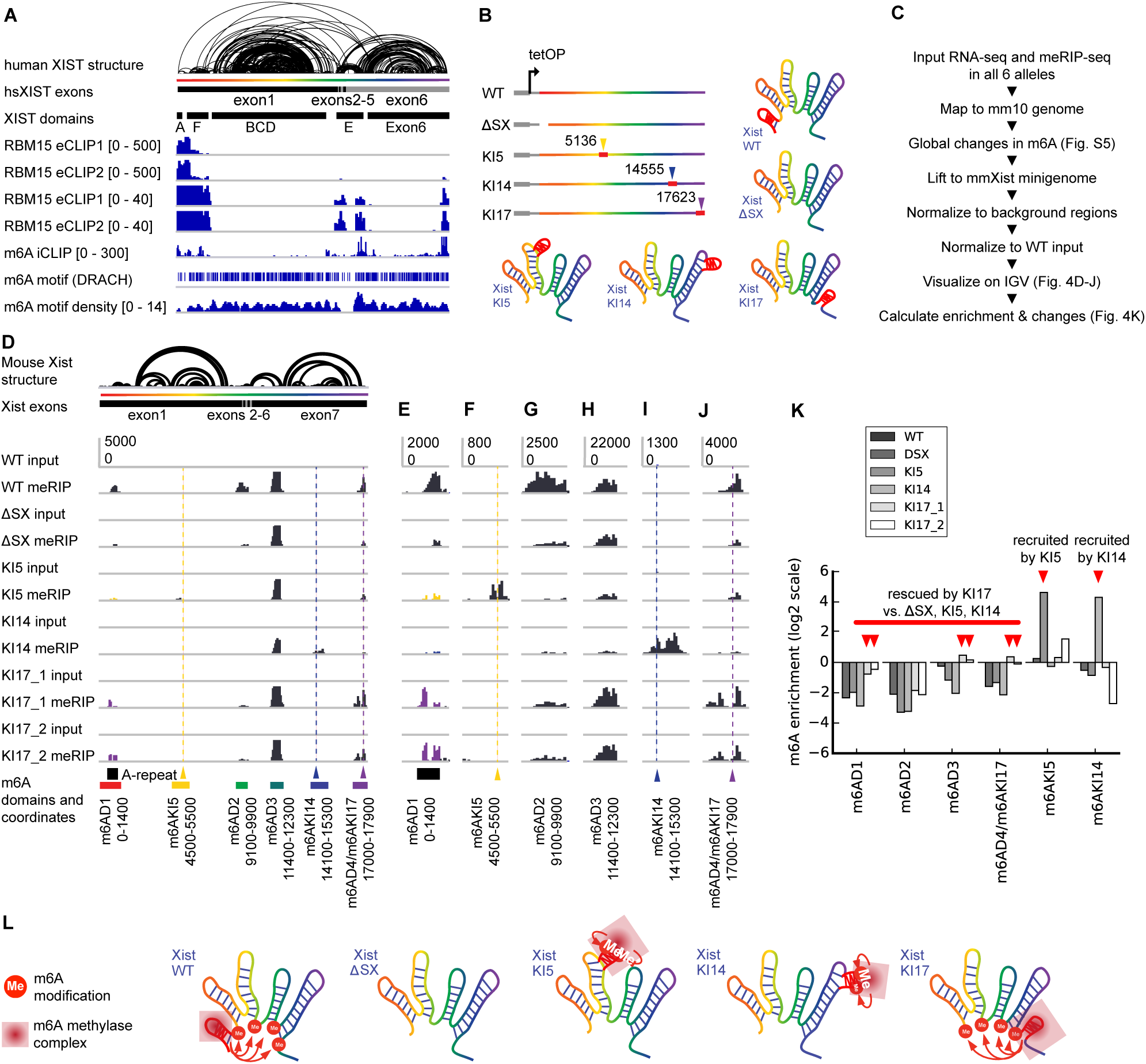
XIST RNA structure determines m^6^A modification patterns. (A) m^6^A modfications on the human mature XIST RNA. The RBM15 eCLIP tracks from K562 cells were normalized against input in 100nt windows (Van Nostrand et al., 2016). Then the enrichment on XIST was plotted in two scales: 0-500 and 0-40, to highlight the differences in binding to the 5’ end and the other regions. The HEK293 cell m^6^A iCLIP track was from (Linder et al., 2015). The m^6^A motif (DRACH) density was calculated in 300nt windows and 50nt steps. (B) Gene structure for the alleles. WT and ΔSX (deletion of ∼900bp in the 5’ end of Xist gene) alleles were from (Wutz et al., 2002), under the control of tetracycline inducible promoter. The A-repeat relocation alleles KI5, KI14, KI17 were derived from ΔSX by insertion of the A-repeat in the indicated locations. Two clones were analyzed for KI17. (C) The pipeline for the meRIP-seq analysis. Global analysis of changes in m^6^A modifications was performed on data mapped to the mm10 genome, while targeted analysis of modifications was performed on data converted to the mature mouse Xist transcript (long isoform, with 7 exons). (D) m^6^A modification sites are changed after relocating the A-repeat domain. The mouse Xist RNA PARIS structure model is the same as in Figure 3A. The m^6^A domains are labeled under the genome browser tracks. The Y-axis is the same in each track. All data including the ones with A-repeat insertion at other locations, were mapped to the same wildtype Xist mature RNA. (E-J). Zoom-in view of all the m^6^A domains. Location of the original A repeat is indicated in panel E. The A-repeat knockin locations are indicated in panels F, I and J. Y-axis scales are the same for all tracks in each panel. (K) Quantification of the changes of m^6^A modification relative to wildtype in log scale in the pre-defined m^6^A domains shown in panel D. (L) A model of the role of RNA structures in guiding m^6^A modifications. The A-repeat domain recruits the m^6^A methylase complex to modify sequences that are physically close the domain. The residual modification on Xist after A-repeat deletion was due to its intrinsic ability to recruit m^6^A methylase complex (see the ΔSX tracks in panels D-J). Relocation of the A-repeat to the inside of the large domains (KI5 and KI14) induces local modifications (m6AKI5 and m6AKI14). Relocation of the A-repeat to the end of the transcript (KI17) induces modification in physical proximity (m6AD1, m6AD2, m6AD3 and m6AD4). See also Figure S4.

To test these hypotheses, we moved the A-repeat sequence to other locations along the Xist RNA by genome editing, and then tested the modification patterns using m^6^A-RIP-seq. We used the J1 male mouse ES cells with an insertion of doxycyclin inducible promoter (here designated as wildtype, or WT), and a derived cell line where the A-repeat region was deleted (ΔSX, removing about 900nt from the beginning of the transcript) (Wutz et al., 2002) (**Figure 4B**). Prolonged induction of WT Xist expression would lead to cell death due to silencing of most genes on the sole X chromosome in these cells, while induction of the ΔSX line did not lead to cell death because absence of the A-repeat region abrogates gene silencing (Wutz et al., 2002). We moved the 440nt A-repeat (less than the deleted region, which is larger than the A-repeat alone) to 3 locations, one in the middle of the large BCD domain (knockin at 5136bp, or KI5), one in the middle of the Exon6 domain (knockin at 14555bp, KI14) and one at the end of the transcript (knockin at 17623bp, KI17) (**Figure S4A**). Relocation to the middle of the large BCD and Exon6 domains is likely to induce local modifications near the insertion sites, while relocation to the end of the transcript is likely to induce modifications in physical proximity.

We generated four isogenic A-repeat insertion lines, one for KI5, one for KI14 and two for KI17 using CRISPR/Cas9 mediated gene editing (**Figure S4**). Then we performed m^6^A-RIP-seq on all 6 cell lines (**Figure 4B**). We measured global changes in m^6^A modifications using m6Aviewer (**Figure S4**), as well as changes on the XIST RNA alone in custom defined regions (**Figure 4C, D**). Four primary m^6^A domains were defined based on proximity: one surrounding the A-repeat region (m6AD1), two around the E domain (m6AD2 and D3), and one at the end of the transcript (m6AD4). In addition we also quantified the modification near the insertion sites (m6AKI5 and m6AKI14). Wildtype mouse Xist harbors m^6^A modification sites in a pattern very similar to the human XIST (compare **Figures 4A** and **D**), and these modification sites are located at the feet of the large RNA domains, whereas the internal regions are almost completely depleted of m^6^A modifications, despite the presence of m^6^A motifs. Removal of the A-repeat greatly reduced m^6^A modification along the entire Xist RNA except for m6AD3, suggesting that the A-repeat is largely required for modifications at distant regions (**Figure 4D-K**). The residual modifications are likely due to the inherent ability of these regions to recruit the m^6^A methylase independent of the A-repeat region.

Insertion of the A-repeat in the middle of the large BCD domain induces modifications near the insertion site (**Figure 4F**), but did not change modification levels at other locations (**Figure 4E, G-J**). Similarly, insertion in the middle of the large Exon6 domain induces local modifications without affecting other regions (**Figure 4I**, compared to panels **4E-H** and **J**). The modifications at these two insertion sites, KI5 and KI14 suggest that the sequences in the middle of the large domains are indeed receptive to modifications, and their lack of modifications in the WT suggest exclusion of the methylase complex (we cannot analyze the exact modified residues due to the lower resolution of RIP-seq). Insertion at the end of the transcript (KI17) led to increases in modifications at all four primary m^6^A domains, m6AD1-4, without affecting the internal regions of BCD and Exon6 domains (m6AKI5 and m6AKI14). The 5’end of the transcript is also modified at higher levels upon A-repeat insertion at the end of the transcript. These data support both hypotheses that the large domains are excluded from modifications, and that regions in physical proximity are modified upon folding of the RNA (**Figure 4K, L**). The XIST RNP can be visualized as a splayed-out hand: The A-repeat is the thumb. Moving the A-repeat “thumb” to the tip of any finger locally affects just that finger. Moving the thumb on the contralateral side of RNA “hand” restores spatial proximity and m6A modification to the base of the RNP hand (**Figure 4L**).

Quantitative RT-PCR analysis showed that each of the KI alleles accumulates lower Xist RNA level than wild type and is unable to induce silencing of X-linked genes (**Figure S5**). Both the lower Xist RNA level and altered RNP architecture may contribute to abrogated Xist function, which will be addressed in future studies.

### Spatial separation XIST RNP functions in binding chromatin and nuclear lamina

During XCI, the HNRNPU family proteins and CIZ1 tether the XIST RNP to the inactive X chromosome, while the LBR protein tethers the XIST RNA to the nuclear lamina (Hasegawa et al., 2010; Kolpa et al., 2016; Ridings-Figueroa et al., 2017; Sakaguchi et al., 2016; Sunwoo et al., 2017). Together, the XIST RNP complex acts as a bridge to bring the X chromosome to the nuclear periphery for remodeling and silencing. To understand how the multiple XIST-associated proteins coordinate the localization of Xi and silencing, we used PIRCh-seq, Purifying Interacting RNAs on Chromatin by sequencing (Fang et al., 2019), to identify regions of RNA that associate with chromatin (**Figure 5A**) and analyzed the binding sites of these proteins on XIST (**Figure 5C**).

**Figure 5.**
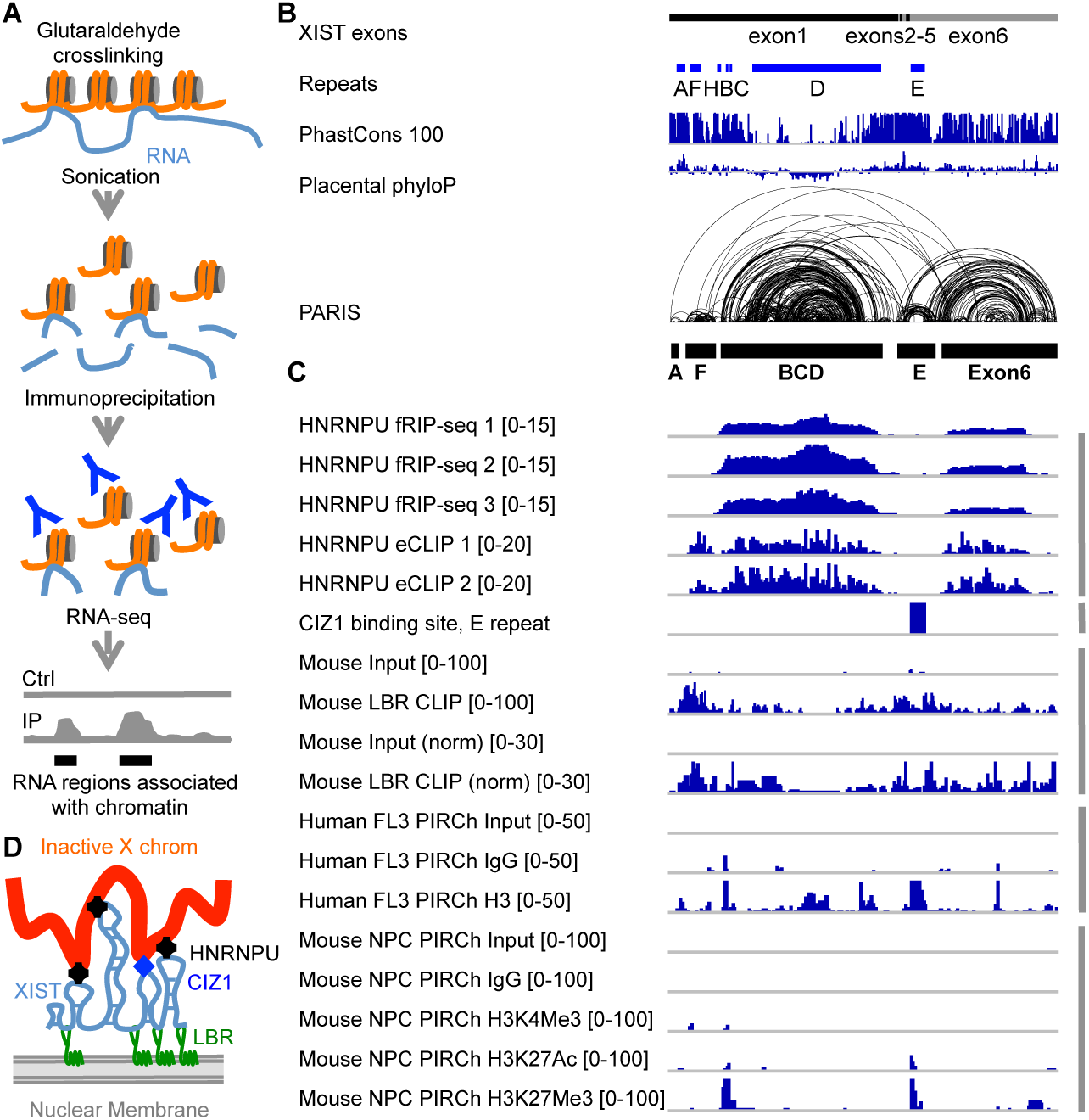
Spatial separation of XIST RNP functions in binding chromatin and nuclear lamina. (A) Schematic diagram of the PIRCh method. Orange lines: genomic DNA, light blue lines: RNA. The Y shape represents antibodies against histones. Ctrl: control, IP: Immunoprecipitation. (B) Annotation of the human XIST RNA. XIST exons and repeats, phylogenetic conservation (PhastCons 100 and Placental PhyloP from UCSC), and PARIS data in HEK293 cells are shown, same as Figure 1D. (C) HNRNPU fRIP-seq (Hendrickson et al. 2016) and eCLIP data in human HEK293 cells (Van Nostrand et al. 2016) were normalized against their input controls in 100nt binds. The mouse CIZ1 binding site track was made based on Ridings-Figueroa et al. 2017, Sunwoo et al. 2017, showing the enrichment of CIZ1 binding on the E-repeat. The LBR raw data and normalized enrichment ratios (in 100nt bins) were from mouse ES cells (Chen et al. 2016), and then lifted to human XIST coordinates. Human and mouse PIRCh data were all normalized against their own controls, respectively in 100nt bins. The mouse PIRCh data were lifted to human XIST coordinates. (D) Model for the spatial separation of XIST RNP functions in binding the inactive X chromosome and the nuclear lamina, color coded the same way as panel (A).

**Figure 6.**
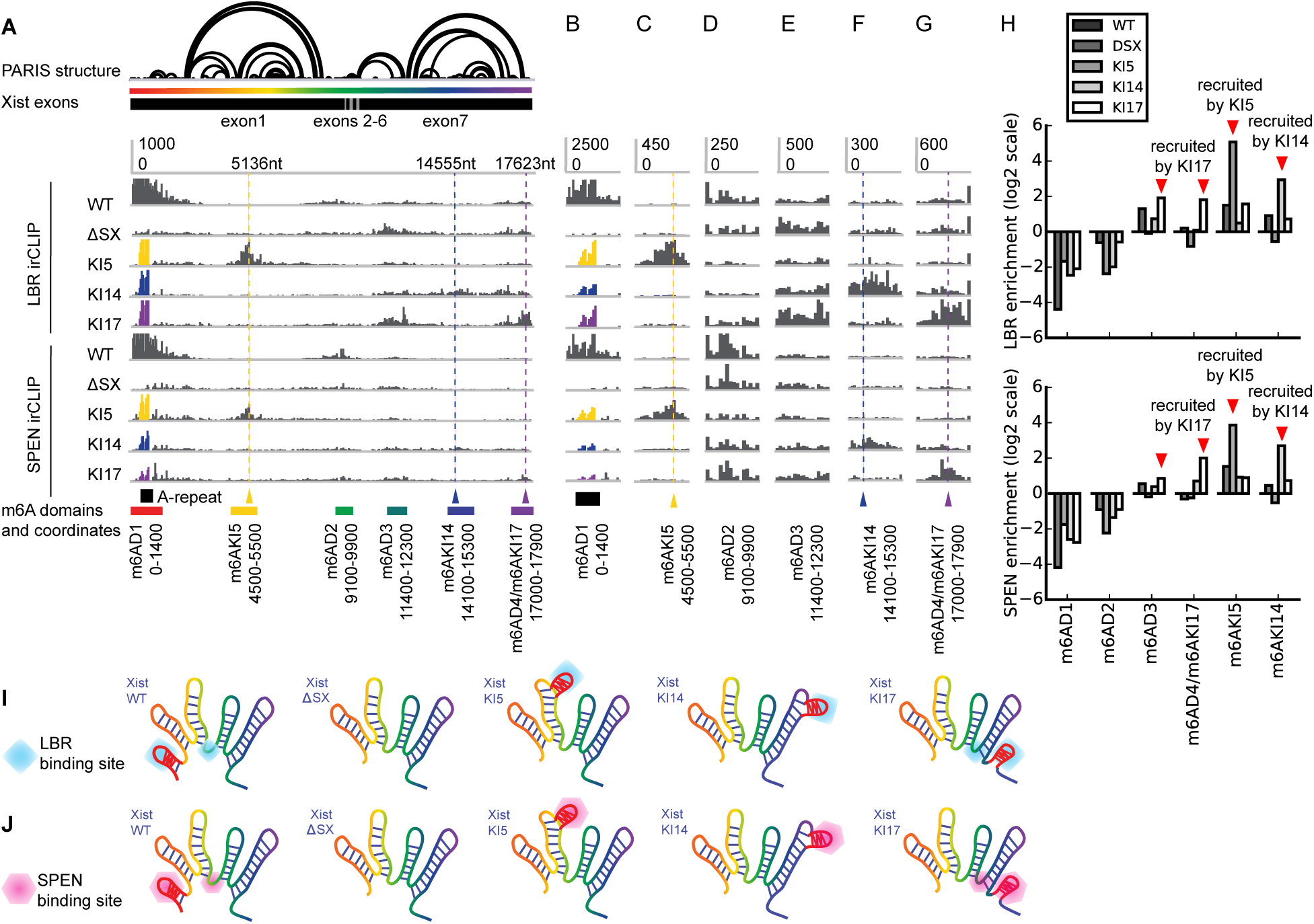
XIST architecture regulates protein-binding specificity. (A) irCLIP of LBR and SPEN in mES cells expressing XIST alleles with relocated A-repeat. The mouse Xist RNA PARIS structure model is the same as in Figure 3A. The m^6^A domains are labeled under the genome browser tracks. The Y-axis is the same in each track. All data including the ones with A-repeat insertion at other locations, were mapped to the same mature wildtype Xist RNA. (B-G) Zoom-in view of all the domains as defined for m^6^A. Location of the original A repeat is indicated in panel B. The A-repeat knockin locations are indicated in panels C, F and G. Y-axis scales are the same for all tracks in each panel. (H) Quantification of the changes of protein binding relative to wildtype in log scale in the pre-defined m^6^A domains shown in panel A. (I-J) Models of the role of RNA structures in guiding protein binding. The A-repeat domain recruits its associated proteins to sequences that are physically close to the domain. The residual binding on Xist after A-repeat deletion was due to its intrinsic ability to recruit proteins (see the ΔSX tracks in panels D-J). Relocation of the A-repeat to the inside of the large domains (KI5 and KI14) induces local binding (m6AKI5 and m6AKI14). Relocation of the A-repeat to the end of the transcript (KI17) binding in physical proximity (m6AD3 and m6AD4). See Figure S4.

fRIP-seq and eCLIP experiments showed that the HNRNPU, CIZ1 (based on PCR) and LBR binding sites are distributed along the XIST RNA (**Figure 1H** and **2**, summarized in **Figure 5C**). In particular, HNRNPU bind the bodies of the large domains, while LBR is enriched on the A-repeat domain and the feet of the larger domains (**Figure 5C**). We hypothesize that XIST regions that are bound by HNRNPU would be more tightly associated with the chromatin, and such regions can be crosslinked to chromatin using glutaraldehyde, a non-specific and highly efficient crosslinker of macromolecules that contain nucleophilic groups like primary amines. After crosslinking, we used sonication to fragment chromatin to small pieces and enriched chromatin-associated RNA using antibodies for histones. The purified RNA were then sequenced to determine relative enrichment (**Figure 5A**). Interestingly, we found that the chromatin-associated regions are primarily in the large domains associated with HNRNPU and CIZ1 (**Figure 5C**), confirming the spatial separation of XIST domains in binding chromatin and nuclear lamina. The enrichment of the C-repeat region by PIRCh is consistent with previous report that showed a role of the C-repeat in chromatin binding (Sarma et al., 2010).

In previous studies, it was noted that LBR binds 3 discrete regions in mouse Xist (around A-repeat domain and flanking the E domain) (Chen et al., 2016). In addition, it was found that deletion of the A-repeat region reduced LBR binding to the latter 2 sites, suggesting cooperative binding. These data also suggest that the sequence motifs alone are not sufficient for LBR binding. In light of the XIST structure model we built, it became clear that the cooperative binding of LBR to 3 distant locations would be mediated by the physical proximity of the folded XIST RNA. Here, the A-repeat domain likely serves to bring LBR to the other locations in physical proximity. We hypothesize that the physical proximity of the A-repeat domain to the feet of the other domains is required for cooperative LBR binding to XIST.

To test the role of A-repeat as a nucleation center, we performed irCLIP on mouse ES cells expressing the A-repeat relocation alleles (**Figure 6A**). Deletion of the A-repeat (ΔSX) reduced LBR binding across the transcript. Insertions of the 440nt A-repeat sequence at 5kb and 14kb not only resulted in the binding at transplanted A-repeats, but also at sequences near the insertion sites, suggesting that the A-repeat is able to recruit protein binding to proximal regions (**Figure 6A-H**). More interestingly, the A-repeat insertion at the end of the transcript (KI17) induced binding to both the 3’ end of the transcript and to m6AD3, which are not contiguous in sequence but are in spatial proximity. Together these data demonstrated that sequence alone was insufficient for protein binding, A-repeat serves as a nucleation center, and the folding of the XIST RNA serves as a conduit for recruiting protein binding to physically close regions (**Figure 6I**). We also performed irCLIP on SPEN, which serves as a bridge to bring the HDAC complex to Xist (Lu et al., 2017). Again, we found that the insertions in the middle of the large domains (KI5 and KI14) resulted in local spreading of SPEN near the A-repeat insertion sites, similar to what we observed for m^6^A modification and LBR binding. The KI17 allele showed spreading around the 17kb insertion site and modest enhancement of binding at m6AD3. Most distant sites such as m6AD1 and m6AD2 were not affected.

## DISCUSSION

### An integrated approach for analyzing large RNP complexes

Long RNA molecules, including the protein-coding mRNAs and lncRNAs, make up the majority of the transcriptome. The dynamic interactions and structures of these RNAs and their protein partners are essential for the exquisite control of gene expression, yet pose major challenges for structure and function analysis. Most of these large RNP complexes are heterogeneous and contain many weak interactions and therefore cannot survive the harsh conditions of purifications for in vitro analysis by crystallography, NMR and cryo-EM. Building upon the PARIS method (Lu et al., 2016), we integrated multiple approaches for the comprehensive characterization of large RNP complexes, including nucleotide flexibility measurements (e.g. icSHAPE) (Spitale et al., 2015), phylogenetic conservation of structures, crosslink and proximity ligation of protein bound RNA structures (e.g. chimeric reads in fRIP-seq), and unsupervised clustering and PCA analysis of RBP binding profiles on RNAs. These orthogonal approaches reveal the organization principles for large RNPs, and their associated functions, from the base pair level, to the domain level (hundreds to thousands of nucleotides). Proximity ligation is the basis of PARIS, and chimeric reads between microRNAs and mRNAs have previously been used to pinpoint microRNA targets (Grosswendt et al., 2014; Helwak et al., 2013). In this work, the crosslink and proximity ligation principle that has been successfully employed in the analysis of chromatin structures and RNA interactions and structures, can be extended to the analysis of other RBPs on any RNA of interest, as long as these proteins are crosslinkable. The application of these methods will be instrumental in the analysis of other RNP complexes. More importantly, the discovery of compact RNP domains set the stage for focused in vitro studies of these domains through purification reconstitution, and structure analysis using physical methods (Lu and Chang, 2018).

### A comprehensive structure-interaction model of the XIST RNP complex

Multiple previous studies have analyzed the XIST RNA, either in part or in its entirety, using various methods (Duszczyk et al., 2011; Fang et al., 2015; Liu et al., 2017; Maenner et al., 2010; Metkar et al., 2018; Smola et al., 2016). Duszczyk et al. determined the in vitro solution structure of a partial A-repeat unit (14nt out of the 24nt unit) using NMR, and found a stable stemloop structure (Duszczyk et al., 2011). Two studies used chemical probing to measure nucleotide flexibility of the in vitro transcribed A-repeat region and found a number of inter-repeat and intra-repeat duplexes (Liu et al., 2017; Maenner et al., 2010). These conflicting in vitro models have not been reconciled. Two additional studies used chemical probing in living cells to analyze XIST RNA structures (Fang et al., 2015; Smola et al., 2016), however, these models may have limitations because, (1) chemical probing reports whether each nucleotide is base paired or constrained by protein binding, and does not directly capture the base pairing relationship, and (2) the secondary structure modeling is based on the faulty assumption that only one stable conformation exists, and thus misses alternative conformations and long range structures (Eddy, 2004; Lu and Chang, 2016).

Using a combination of five orthogonal methods, we have built a comprehensive model of the XIST RNP complex. In our previous study, we have applied PARIS, icSHAPE and phylogenetic analysis to determine the overall structure of the XIST RNA. In the current study, we incorporated systematic analysis of RNA-protein interactions data based on fRIP-seq and eCLIP and also examined the proximally ligated reads from RNA-protein interactions. Recently, Moore and colleagues used formaldehyde to crosslink the exon junction complex to RNAs and found long-distance RNA structures in XIST similar to the PARIS-derived modular domains (Metkar et al., 2018). These studies firmly established the modular architecture of the entire RNP complex. Sequence inside the module are more likely to basepair with each other and also interact with similar RBPs, while sequences outside of the module may be excluded for interactions.

The discovery of this modular topology raises many questions about the formation and function of XIST structures. While we have found a critical role of the XIST structure in determining RBP specificity, it is unclear how the long-range structures form in the first place. RNA structure formation is primarily driven by the stacking of base pairs. As RNA is transcribed, local structures can quickly form low energy conformations, and the co-transcriptional folding process is integral to the function of riboswitches in bacteria (Frieda and Block, 2012). Thus mechanisms must exist to counter the tendency of local structure formation to allow long-range structures in XIST to form. Given the high abundance of many hnRNP proteins in the nucleus and their interactions with XIST (e.g. HNRNPK and HNRNPU), it is likely that the co-transcriptional binding of hnRNPs would compete with base pairing and contribute to the formation of observed high level architecture. Certain hnRNP proteins possess RNA annealing activities, and as a result may affect the observed structures by remodeling of co-transcriptionally formed duplexes (Herschlag, 1995; Portman and Dreyfuss, 1994). Numerous RNA helicases associate with the XIST RNA and likely play a role in remodeling XIST structure (Chu et al., 2015).

Multivalent interactions underlie phase separation, a general mechanism of organization in cells. Recently it was reported that the XIST RNP complex forms a distinct liquid phase in cells, probably due to the repetitive sequences in XIST and the intrinsically disordered regions in XIST-associated proteins (Cerase et al., 2018). Our analysis showed that XIST associated proteins bind XIST in clustered patterns, supporting the multivalent interactions. Therefore, the phase separation process might contribute to the high affinity and specificity of XIST RNP formation.

### The A-repeat domain as a nucleation center for multiple XIST functions

The repetitive nature of the A-repeat has made it a challenging target for structure analysis. As discussed above, several studies have reported structure models for the A-repeat region using chemical probing methods, leading to conflicting models (Lu et al., 2017; Maenner et al., 2010). We have used several methods to establish a stochastic inter-repeat duplex model (Lu et al., 2016). Using CLIP and gel shift assays we found that the A-repeat region forms a multivalent platform to bind the SPEN adapter protein. Importantly, our model has been confirmed by a more recent rigorous phylogenetic analysis of noncoding RNA structure conservation (Rivas et al., 2016). In the current study, we further extended the model of the essential A-repeat domain by identifying additional potential interactions, including RBM15/RBM15, SRSF1, U2AF1, and LBR. This multitude of high affinity A-repeat-associated proteins suggests that the A-repeat serves as a nucleation center for recruiting XIST binding proteins. Using the A-repeat relocation XIST alleles and CLIP experiments, we showed that the A-repeat is indeed sufficient in spreading physically local RBP binding and m^6^A modifications (**Figures 4** and **6**). The nucleation function of A-repeat together with the topology of the entire XIST RNA are responsible for generating the unique patterns of protein binding, as well as the functions associated with these proteins. The multiple proteins binding to the A-repeat create a crowded environment, and it remains to be determined how these proteins are assembled in spatial and temporal specific manner. For example, the splicing factors are required early on for proper XIST maturation, while the effectors of XIST functions may bind XIST later. It has been shown that XIST interactions with proteins changes during stem cell differentiation (Chu et al., 2015). It is conceivable that a dynamic process of XIST RNP assembly coordinates its functions. In addition, other ways of organizing the functions are also possible.

### Spatial separation and coordination of XIST functions by the modular architecture

XCI is a complex process and XIST coordinate multiple steps in this process. We have found that the XIST-associated functions are spatially separated on the RNA structural scaffold. For instance, the A-repeat domain together with the physically close regions of the RNA located at the feet of the other domains associate with proteins involved in transcriptional silencing, m^6^A modification, splicing regulation, DNA methylation and nuclear lamina attachment. The body of the large BCD, E and exon6 domains binds HNRNPU family proteins and CIZ1, which then tethers XIST to the inactive X chromosome. The mechanisms that coordinate other functions in XIST remain to be discovered. The discovery of the modular domain architecture provides a framework for future analysis of other functions coordinated by XIST.

Previous studies on the yeast 1.2kb long telomerase RNA revealed an RNA scaffold that contains several essential protein binding sites (Zappulla and Cech, 2004). The arms of the RNA scaffold can be relocated without affecting their functions. Thus the individual domains in TERC RNA are like words on a billboard; their presence rather than exact order convey most of the message. Here our studies revealed a more complex picture for the flexible domain architecture of XIST: the scaffold serves to insulate certain regions and bridge other regions. The role of structure-induced proximity in guiding RBP binding and m^6^A modification is similar to the concept of chromatin conformations guiding gene expression regulations. Chromatin structures can either insulate regions from surrounding epigenetic environment, or induce spatial proximity to bring regulatory elements to gene promoters. For example, the structure of the X chromosome guides the spreading of XIST to spatially close sites (Engreitz et al., 2013). The specific order of domains in XIST suggests the existence of grammar rules in lncRNA function. The nature of this grammar, whether for stable RNP assembly or RNP function remains to be determined.

## ACKNOWLEDGEMENT

We thank CK Chen and M Guttman for help with LBR and SPEN CLIP. Supported by NIH R01-HG004361 and RM1-HG007735 (H.Y.C), King Abdulaziz University (H.C., H.Y.C). Z.L. was a Layton Family Fellow of the Damon Runyon-Sohn Foundation Pediatric Cancer Fellowship Award (DRSG-14-15), and supported by Stanford Jump Start Award of Excellence in Postdoctoral Research and the Pathway to Independence Award from NHGRI (1R00HG009662). We also acknowledge the USC Norris Comprehensive Cancer Center (P30CA014089) for their support of our research. H.Y.C. is an Investigator of the Howard Hughes Medical Institute.

## AUTHOR CONTRIBUTIONS

Conceptualization, Z.L. and H.Y.C.; Methodology, Z.L., J.K.G, F.L. and Q.M.; Investigation, Z.L., J.K.G, D.R.D., B.Z., F.L., H.C., Q.M., and P.A.K.; Data Analysis: Z.L., Y.W.; Writing, Z.L. and H.Y.C.; Funding Acquisition, Z.L. and H.Y.C.; Resources, H.Y.C.; H.C. Supervision, Z.L. and H.Y.C.

---

**Figure S1.**
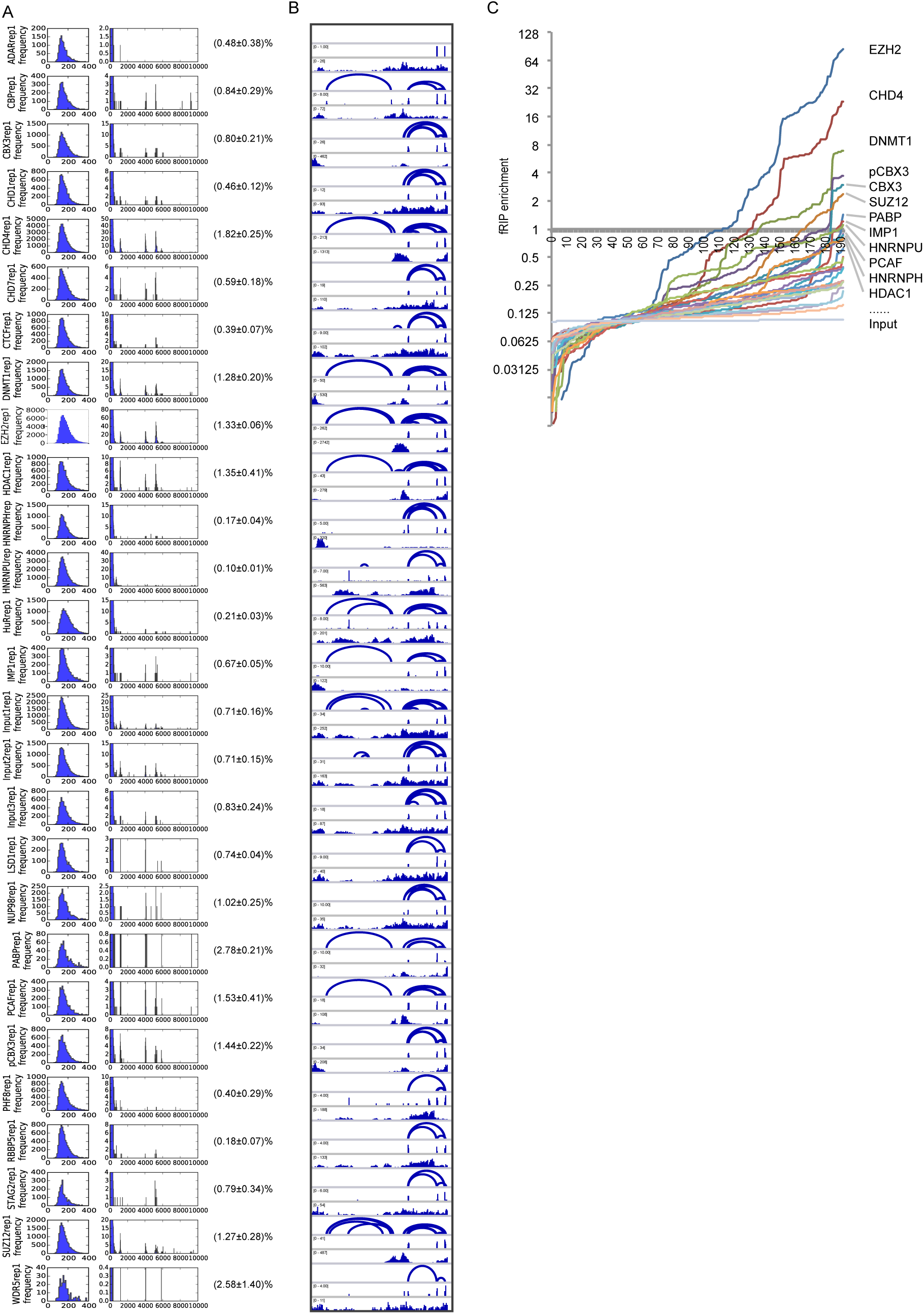
fRIP-seq data for all proteins. Related to Figure 1. (A) Distance distribution of the short-distance and long-distance read pairs from paired-end fRIP-seq data. See Figure 1 for description of the panels. Here one replicate is selected from each control or fRIP-seq experiment. The percentage of long distance read pairs in all read pairs, calculated as average ± standard deviation are shown to the right of the panel (n=2 for EZH2 and STAG2, n=3 for the others). (B) IGV genome browser tracks for all the fRIP-seq data, showing one replicate from each experiment. For each sample, the 1st track, geometric_long_arc.bed, presents the connections of the two sequencing tags in each paired-end read; the second track, geometric_long.bedgraph, presents the depth of long-distance read pairs (>1000nt); the third track, geometric.bedgraph, presents the depth of all reads mapped to XIST. (C) Enrichment of proteins on XIST in 100nt bins, sorted in ascending order in log scale. The top enriched proteins and input are labeled. The X-axis indicates the ranks of the 193 bins.

**Figure S2.**
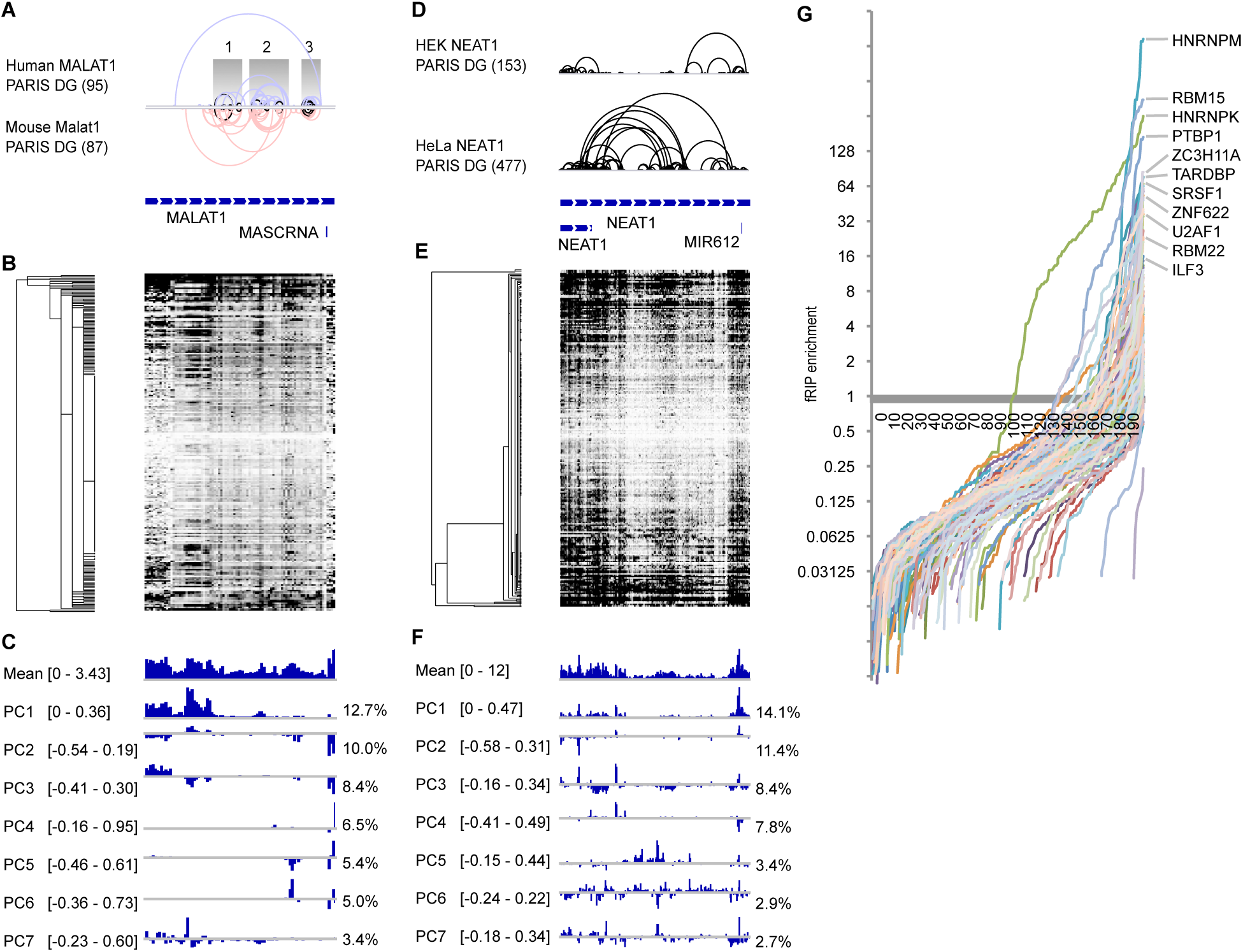
eCLIP analysis of protein binding on lncRNAs MALAT1 and NEAT1. Related to Figure 2. (A, D) PARIS derived structure models and gene models for MALAT1 (A) and NEAT1 (D) from (Lu et al., 2016). (B, E) Clustering of protein enrichment profiles in 100nt windows for all 242 samples of the 121 proteins, two biological replicates each, for MALAT1 (B) and NEAT1 (E). (C, F) PCA analysis of all eCLIP data in 100nt windows for MALAT1 (B) and NEAT1 (E). The mean and first 7 principal components are displayed together with percentage of variation explained by each component on the right. Together, these principal components explain 51% of total variation in each lncRNA.

**Figure S3.**
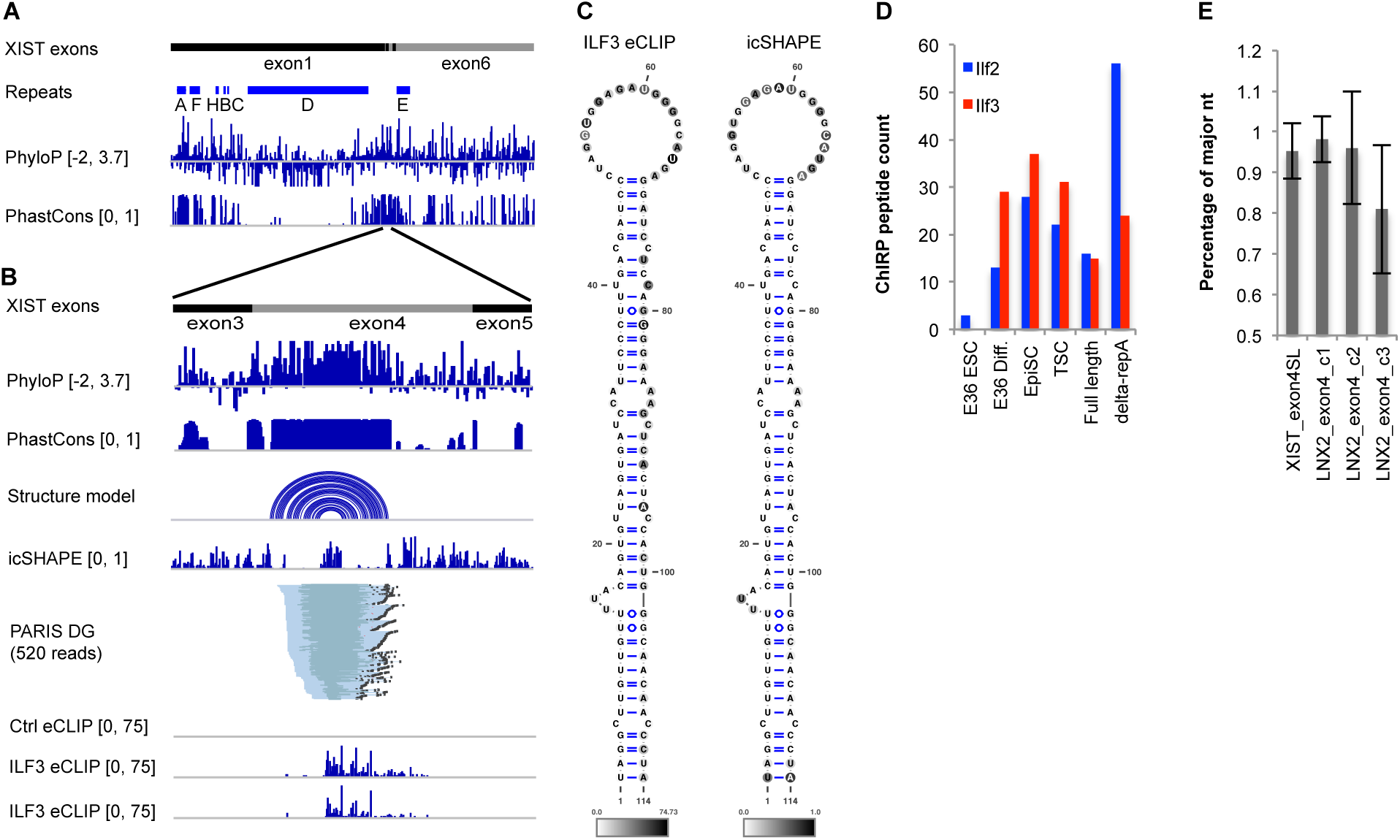
Structure and interaction model of the conserved Exon 4 stemloop. (A) Annotation of the human XIST mature transcript. The 6 exons in human XIST were concatenated, with the introns removed. The repeat elements were annotated based on (Elisaphenko et al., 2008). The placental mammals phyloP and 100 vertebrates PhastCons were from UCSC. (B) XIST exon 4 forms a conserved stemloop structure that interacts ILF3. The icSHAPE and PARIS data were from human HEK293T cells (Lu et al., 2016), while the ILF3 eCLIP data were from human K562 cells (Van Nostrand et al., 2016). ILF3 binds the loop and right side of the stem. Note, the stem-loop region in exon4 is highly conserved, while the rest of exon4 is not. (C) The icSHAPE and ILF3 eCLIP data are plotted on the human XIST exon 4 stemloop structure. (D) Both ILF3 and ILF2 were previously identified as XIST interacters (Chu et al., 2015). Peptide counts from XIST ChIRP-MS were plotted against cell lines used that indicate different stages in stem cell differentiation. (E) The conservation of the stemloop region in exon4 in eutherian mammals as compared to the ancestral LNX2 exon4 region. Percentage of the dominating nucleotide at each position was calculated. LNX2 third codon position is less conserved consistent with the wobble position.

**Figure S4.**
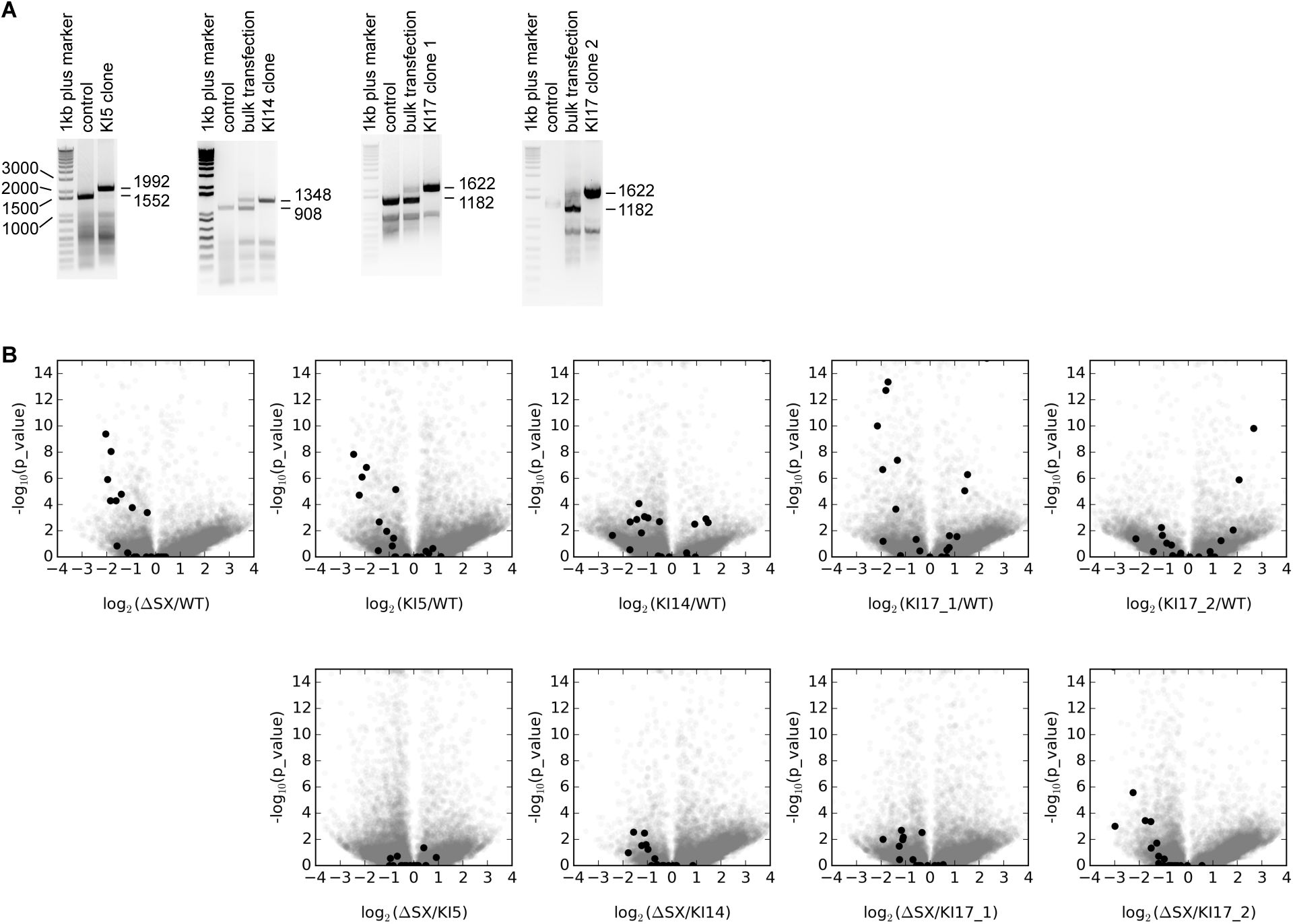
Relocation of the A domain alters m^6^A modification patterns. Related to Figure 4. (A) Genotyping PCR for the four clones picked for A-repeat relocation. Control samples are the starting ΔSX cell line (A repeat deletion). PCR for bulk transfections were performed after CRISPR editing and before picking clones. Molecular size markers are in base pairs. (B) Global changes of m^6^A modifications after relocation of the A-repeat domain. m^6^A modification sites were identified using the m6aViewer software (version 1.6.1) with the default parameters and negative strand bam files (half the data). Differences were visualized in volcano plots in the selected pairs of comparisons. Log scale fold changes (LFC) and negative log scale p values were used as the x-axis and y-axis, respectively.

**Figure S5.**
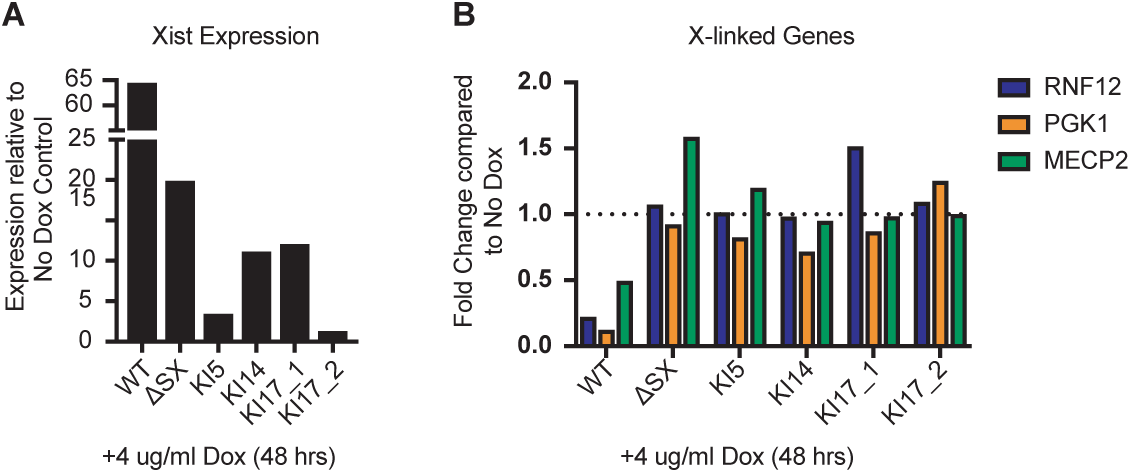
Analysis of X chromosome inactivation after relocation of the A-repeat. Related to Figure 4. (A). A-repeat relocation cell lines were induced to express Xist with doxycycline treatment and Xist levels were measured using qRT-PCR. (B). A-repeat relocation cell lines were induced to express Xist. Then the expression of X-linked genes were measured using qRT-PCR.

**Table S1. Enrichment of RBPs on XIST, and comparison to ChIRP-MS. Related to** **Figure 2.** The 121 RBPs studied by eCLIP were analyzed using window-based normalization against size-matched controls (See Supplementary Methods for details). The entire 19278nt human mature XIST is divided into 193 windows, each 100nt, and the enrichment values were ranked from low to high for each RBP. The geometric mean of enrichment ratios for the two biological replicates for each RBP were used for the ranking. The 25^th^ percentile of non-zero enrichment values in each RBP was set to 0.1. Then the highest enrichment bin was used to rank all RBPs. Out of the 81 proteins previously detected by Xist ChIRP in mouse cells (Chu et al., 2015), 27 of them are included in the 121 eCLIP dataset, and highlighted.

## STAR METHODS

Detailed methods are provided in the online version of this paper and include the following:

### KEY RESOURCES TABLE

| REAGENT or RESOURCE | SOURCE | IDENTIFIER |
| --- | --- | --- |
| **Antibodies** |  |  |
| Anti-N6-methyladenosine (m6A) antibody | Millipore Sigma | ABE572 |
| SPEN antibody | Bethyl Laboratories | A301-119A |
| SPEN antibody | Abcam | Ab72266 |
| RBM15 antibody | Protein Tech | 10587-1-AP |
| RBM15B antibody | Protein Tech | 22249-1-AP |
| LBR antibody | Protein Tech | 12398-1-AP |
| LBR antibody | Guttman laboratory |  |
| **Chemicals, Peptides, and Recombinant Proteins** |  |  |
| 4'-Aminomethyltrioxsalen hydrochloride | Sigma-Aldrich | Cat# A4330 |
| Cas9 Protein with NLS | PNA Bio | Cat# CP01 |
| Doxycycline | Sigma-Aldrich | Cat# D9891 |
| **Critical Commercial Assays** |  |  |
| SMARTer Stranded Total RNA-Seq Kit v2 - Pico Input Mammalian | Clontech | Cat# 634411 |
| Zero Blunt TOPO PCR Cloning Kit, One Shot | Thermo Fisher | Cat# K2875 |
| **Deposited Data** |  |  |
| fRIP-seq | (Hendrickson et al., 2016) | GEO: GSE67963 |
| eCLIP | (Van Nostrand et al., 2016) | https://www.encodeproject.org/ |
| LBR CLIP | (Chen et al., 2016) | GEO: GSE86250 |
| m6A iCLIP | (Linder et al., 2015) | GEO: GSE63753 |
| Human HEK293 cell PARIS data | (Lu et al., 2016) | GEO: GSE74353 |
| Human and mouse PIRCh | (Fang et al., 2019) | GEO: GSE119006 |
| Mouse ES cell PARIS | This study | GEO: GSE126716 |
| Mouse ES cell meRIP-seq and irCLIP | This study | GEO: GSE126715 |
| **Experimental Models: Cell Lines** |  |  |
| HATX mES cell line | Anton Wutz | (Monfort et al., 2015) |
| TXY mES cell line | Edith Heard | (Wutz et al., 2002) |
| TXY:ΔSX mES cell line | Edith Heard | (Wutz et al., 2002) |
| TXY:KI5, TXY:KI14, TXY:KI17_1, TXY:KI17_2 mES lines | This paper |  |
| **Oligonucleotides** |  |  |
| Guide RNA templates | This paper | Details below |
| Cloning and testing primers | This paper | Details below |
| Guide RNA test primers | This paper | Details below |
| **Software and Algorithms** |  |  |
| STAR | (Dobin et al., 2013) | https://github.com/alexdobin/STAR |
| Samtools v1.1 | (Li et al., 2009) | http://samtools.sourceforge.net/ |
| Bedtools v2.22.0 | (Quinlan and Hall, 2010) | https://bedtools.readthedocs.io/ |
| m6aViewer v1.6.1 | (Antanaviciute et al., 2017) | http://dna2.leeds.ac.uk/m6a/ |
| PARIS | (Lu et al., 2016) | https://github.com/qczhang |
| IGV | (Robinson et al., 2011) | http://broadinstitute.org/software/igv |
| Vienna RNA Package | (Lorenz et al., 2011) | https://www.tbi.univie.ac.at/RNA/ |
| Kent Utilities | (Fujita et al., 2011) | https://genome.ucsc.edu/util.html |
| Trimmomatic v0.3.2 | (Bolger et al., 2014) | http://www.usadellab.org/cms/?page=trimmomatic |
| Python | (Van Rossum, 1995) | https://www.python.org/ |
| Cluster | (de Hoon et al., 2004) | http://bonsai.hgc.jp/~mdehoon/software/cluster/software.htm |
| Java Treeview | (Saldanha, 2004) | http://jtreeview.sourceforge.net/ |
| Fastqc | (Andrews, 2010) | https://www.bioinformatics.babraham.ac.uk/projects/fastqc/ |
| Custom scripts | This paper | https://github.com/zhipenglu |

### CONTACT FOR REAGENT AND RESOURCE SHARING

Further information and requests for reagents should be directed to and will be fulfilled by the Lead Contact, Howard Y. Chang.

### EXPERIMENTAL MODEL AND SUBJECT DETAILS

#### mES Cell culture

Male inducible TXY WT, TXY: ΔSX lines (gift from Anton Wutz) (Wutz et al., 2002), and all generated TXY knockin cell line derivatives (TXY:KI5, TXY:KI14, TXY:KI17) were cultured and treated for 48h with 4 ug/ml doxycycline before RNA collection. HATX3 cells (*Xist^TX/TX^ Rosa26^nlsrtTA/nlsrtTA^*) were grown under the same conditions before AMT crosslinking and RNA collection (Monfort et al., 2015). All mES cells were maintained on 0.2% gelatin coated plates at 37C with mES media, which was changed daily: Knockout DMEM + 10% FBS + 1% MEM NEAA + 1% GlutaMax + 1% Pen Strep + 0.2% BME and 0.01% LIF.

### METHOD DETAILS

#### Analysis of fRIP-seq data

To determine the interactions between XIST and associated proteins, fRIP-seq experiments on 24 chromatin-associated and traditional RNA binding proteins in human K562 cells were reanalyzed (GSE67963) (Hendrickson et al., 2016). In the fRIP-seq experiments, formaldehyde crosslinked RNP complexes were sonicated so that the associated RNA fragments are around a few hundred nucleotides, and the sequenced fragments are around 150nt (see Figure 1 and S1 for size distribution). Paired end sequencing was performed on the libraries, 31nt each end. The general pipeline is as follows: Convert paired end reads to gap reads map to hg38 extract reads mapped to XIST map to hsXIST ‘minigenome’ assemble mapped Aligned and Chimeric reads ex tract long pairs make distance distribution, bedgraph and arcs f or both short and long pairs.

1. Download data from sra using the following standard format: /sra/sra-instant/reads/ByRun/sra/{SRR|ERR|DRR}/<FIRST 6 characters of accession>/<ACCESSION>/<ACCESSION>.sra. For example: wget ftp://ftp-trace.ncbi.nih.gov/sra/sra-instant/reads/ByRun/sra/SRR/SRR197/SRR1976881/SRR1976881.sra

2. Convert sra to fastq using fastq-dump in the sra-toolkit, convert from paired end fastq to gapped fastq using the pe2gap.py script, and then combine the multiple fastq files for each sample (for example, concatenate SRR1976598-SRR1976603 to ADARrep1). The pe2gap.py script converts the second half of each 62nt read to its reverse complement. for file in SRR*sra; do (fastq-dump $file &); done for file in *fastq; do (python pe2gap.py 32 $file ${file%fastq}gap.fastq &); done

3. Map the combined fastq files to hg38 with the following specific parameters. The limitOutSJcollapsed option is adjusted to accommodate the large number of ‘splice junctions’ because essentially all the gapped reads are like ‘splice junctions’. The chimSegmentMin option allows chiastic mapping.

for file in *gap.fastq; do (star-static --runMode alignReads --genomeDir hg38/ --readFilesIn $file --outFileNamePrefix

${file%.fastq}_hg38 --outReadsUnmapped Fastq --outFilterMultimapNmax 1 --outSAMattributes All --alignIntronMin 1 --chimSegmentMin 15 --chimJunctionOverhangMin 15 --limitOutSJcollapsed 3000000 --runThreadN 8 &); done

4. Use samtools to convert, sort and index the star output sam files

for file in *gap_hg38Aligned.out.sam; do (samtools view -bS -o ${file%out.sam}bam $file; samtools sort

${file%out.sam}bam ${file%out.sam}_sorted; samtools index ${file%out.sam}_sorted.bam &); done

for file in *gap_hg38Chimeric.out.sam; do (samtools view -bS -o ${file%out.sam}bam $file; samtools sort

${file%out.sam}bam ${file%out.sam}_sorted; samtools index ${file%out.sam}_sorted.bam &); done

5. Convert the reads mapped to XIST in hg38 back to fastq. Note the output redirection is different for the two files.

for file in *gap_hg38Aligned_sorted.bam; do (samtools view $file chrX:73820651-73852753 | awk ’{print "@" $1 "\n" $10 "\n+\n" $11}’ > ${file%Aligned_sorted.bam}XIST.fastq &); done

for file in *gap_hg38Chimeric_sorted.bam; do (samtools view $file chrX:73820651-73852753 | awk ’{print "@" $1 "\n" $10 "\n+\n" $11}’ >> ${file%Chimeric_sorted.bam}XIST.fastq &); done

6. Map XIST reads to hsXIST ‘mini-genome’, which consists of the human mature XIST RNA sequence, with the following specific parameters. See previous publication on how the mini-genome was made (Lu et al., 2016).

for file in *gap_hg38XIST.fastq; do (star-static --runMode alignReads --genomeDir starhsXIST/ --readFilesIn $file -- outFileNamePrefix ${file%hg38XIST.fastq}hsXIST --outReadsUnmapped Fastq --outFilterMultimapNmax 1 -- outSAMattributes All --alignIntronMin 1 --chimSegmentMin 15 --chimJunctionOverhangMin 15 --runThreadN 8 &); done

7. Use samPairingCalling.test.pl (from https://github.com/qczhang/) to convert the chiastic reads to normal gapped reads. This step is only used to combine the two files, not to assemble the duplex groups. The duplex group information is not used in the analysis of the five groups of long-distance reads.

for file in *gap_hsXISTAligned.out.sam; do (perl samPairingCalling.test.pl -i $file -j

${file%Aligned.out.sam}Chimeric.out.junction -s ${file%Aligned.out.sam}Chimeric.out.sam -o

${file%Aligned.out.sam}_geometric -g starhsXIST/hsXIST.fa -z starhsXIST/chrNameLength.txt -a annotations/empty.gtf -t starhsXIST/hsXIST.fa -l 15 -p 2 -c geometric 1>${Aligned.out.sam}_geometric.stdout 2>${Aligned.out.sam}_geometric.log &); done

8. Remove improperly assembled DGs (duplex groups), which are caused by lack of sufficient support reads. for file in *gap_hsXIST_geometricsam; do (grep -v "DG:i:$" $file > ${file%sam}.sam &); done

9. Use readspan.py to extract the long pairs from the *gap_hsXIST_geometric.sam files and plot all the length distributions, output long.sam, shortdist.pdf and longdist.pdf. The cutoff is set at 1000nt to ensure only proximity-ligated fragments are extracted, given the length of average RNA fragments less than 200nt. The pdf files are assembled into multi-panel figures (Figure 1 and S1).

for file in *gap_hsXIST_geometric.sam; do (python readspan.py ${file%gap_hsXIST_geometric.sam} $file

${file%.sam}_long.sam ${file%.sam}_shortdist.pdf ${file%.sam}_longdist.pdf &); done

10. Make bedgraph files for visualization on IGV. The following is the script for automated processing of all samples. These tracks are combined with the arcs produced below (step 11) into multi-panel figures (Figure 1 and S1).

for file in *gap_hsXIST_geometric*.sam; do (samtools view -bS -o ${file%sam}bam $file; samtools sort ${file%sam}bam

${file%.sam}_sorted; samtools index ${file%.sam}_sorted.bam; genomeCoverageBed -ibam ${file%.sam}_sorted.bam -bg

-split -g hsXIST.size > ${file%sam}bedgraph &); done

11. Use cigar2helixbed.py to convert the *gap_hsXIST_geometric_long.sam to bed files to visualize the arcs in the IGV genome browser. See IGV documents for the instructions on the visualization (https://software.broadinstitute.org/software/igv/node/284).

for file in *gap_hsXIST_geometric_long.sam; do (python cigar2helixbed.py $file ${file%.sam}_arc.bed &); done

12. Use the frip_subset_paris.py script to group the long-distance read pairs as follows. Given that only five major groups are discernable, reads are grouped by their anchor locations. The long-distance groups (LGs) are visualized together with the overlapping DGs from PARIS data (from step 13) in Figure 1.

for file in *gap_hsXIST_geometric_long.sam; do (python frip_subset_paris.py $file ${file%.sam}anchors.sam; samtools view -bS -o ${file%.sam}anchors.bam ${file%.sam}group.sam; samtools sort ${file%.sam}anchors.bam

${file%.sam}anchors_sorted; samtools index ${file%.sam}anchors_sorted.bam &); done

13. Use the frip_subset_paris.py script to extract PARIS DGs that are in the same region as the 5 fRIP-seq long-distance groups (LGs) as follows.

for file in AMT_Stress_trim_nodup_starhsXIST_l15p2_geometricNGmin.sam; do (python frip_subset_paris.py $file

${file%.sam}anchors.sam; samtools view -bS -o ${file%.sam}anchors.bam ${file%.sam}group.sam; samtools sort

${file%.sam}anchors.bam ${file%.sam}anchors_sorted; samtools index ${file%.sam}anchors_sorted.bam &); done

14. The 31nt sequencing tags do not represent the actual binding sites of the proteins; instead, the tags need to be extended to the size of the RNA fragments (∼150nt) to reveal the approximate location of the protein binding and crosslinking. To make the extended fRIP-seq profiles for the long-distance pairs, the following script was used:

frip_extend_longpairs.py. The data are visualized in Figure 1 and S1.

python frip_extend_longpairs.py 119 EZH2rep1gap_hsXIST_geometric_longanchors_LG2.sam EZH2rep1gap_hsXIST_geometric_longanchors_LG2extend.sam

15. To normalize the bedgraph files for visualization on IGV, we made one copy of Input1rep1 control for each IP, since the level of enrichment is very different, a single normalization would not work. For example, for HNRNPU we use the first 800nt given the clustered binding in the large BCD and exon6 domains. The count for Input1rep1 is 664, and for HNRNPUrep1 is 66. Therefore the ratio is 0.099. For CBX3, we used the region 2000nt to 10000nt as follows and the normalization factor is 0.298

awk ’($4>2000)&&($4<10000)’ Input1rep1gap_hsXIST_geometric.sam | wc -l

awk ’{print $1 "\t" $2 "\t" $3 "\t" $4*0.298}’ CBX3rep1gap_hsXIST_geometric.bedgraph > CBX3rep1gap_hsXIST_geometric_norm.bedgraph

16. To cluster the profiles of fRIP-seq experiments, the following commands are used. First we made 100nt windows and calculated coverage in 100nt intervals. Then all the files were combined to generate a matrix for all fRIP-seq profiles.

bedtools makewindows -g hsXIST.size -w 100 > hsXIST_100nt.bed

bedtools coverage -split -abam ADARrep1gap_hsXIST_geometric_sorted.bam -b hsXIST_100nt.bed | sort -k2 -n >

ADARrep1gap_hsXIST_geometric_100nt.bed

for file in *gap_hsXIST_geometric_sorted.bam; do (bedtools coverage -split -abam $file -b hsXIST_100nt.bed | sort -k2 -n

| cut -f4 > ${file%sorted.bam}100nt.vector &); done

awk ’{print $1 "_" $2 "_" $3}’ ADARrep1gap_hsXIST_geometric_100nt.bed > frip_gap_hsXIST_geometric_100nt.intervals add a column name to the frip_gap_hsXIST_geometric_100nt.matrix file

for file in *gap_hsXIST_geometric_sorted.bam; do (echo ${file%gap_hsXIST_geometric_sorted.bam} | tr ’\n’ ’\t’ >> frip_gap_hsXIST_geometric_100nt.matrix); done

add a new line to the end of the frip_gap_hsXIST_geometric_100nt.matrix file

paste frip_gap_hsXIST_geometric_100nt.intervals *vector >> frip_gap_hsXIST_geometric_100nt.matrix

17. To normalize the frip_gap_hsXIST_geometric_100nt.matrix file against input controls, we first divided the values of each bin in each sample by the values of the average of Input1rep1-Input1rep3. Then we adjusted the 25th percentile of each sample to 0.1. This step was performed using the script frip_norm.py.

18. After the matrix file was normalized, the clustering was performed using Cluster 3.0 and Java TreeView (de Hoon et al., 2004; Saldanha, 2004). Cluster 3.0 options: Hirarchical, Arrays Cluster, correlation (uncentered), complete linkage. TreeView options as follows. Settings: Pixel settings: Conrast 3.0. Positive black, zero white. Export to postscript, Gene Headers: NAME, Array Headers: Interval, Below tree: yes. Include: Array Tree, x scale 7, y scale 3.5, Border 0.

19. To plot the RIP enrichment of each sample in 100nt bins, replicates for each sample were averaged, sorted in ascending order and then plotted in Excel.

#### Analysis of eCLIP data

The Yeo lab performed large-scale CLIP experiments to determine the transcriptome wide binding sites of over 100 proteins in female K562 cells and male HepG2 cells (Van Nostrand et al., 2016), and reported four proteins that bind XIST specifically (>2 fold enrichment). To determine the interactions between XIST and associated proteins, the eCLIP data were reanalyzed as follows. The total numbers of bigwig files used are 726 and 618 respectively and the files are named as follows K562/HepG2_RBP_0/1/2_neg/pos.bw.

1. Download the bigWig files for all K562 eCLIP data from ENCODE (https://www.encodeproject.org/search/?type=Experimentl) using the following selection criteria: Assay: eCLIP, Experiment status: released, Biosample type: immortalized cell line, Life stage: adult, Available data: bigWig, and the following standard command: xargs -n 1 curl -O -L < files.txt. The downloaded bigWig files were renamed using the metadata.tsv file linked within the files.txt file.

For data that were mapped to hg19, they were converted to hg38 using eclip_rename.py as follows. for file in *hg19.bw; do (bigWigToBedGraph $file ${file%bw}bedgraph &); done

for file in *hg19.bedgraph; do (liftOver $file ∼/annotations/hg19ToHg38.over.chain ${file/_hg19/} ${file}unmapped &); done for file in *neg.bedgraph *pos.bedgraph; do (sort -k1,1 -k2,2n $file > ${file%.bedgraph}sorted.bedgraph &); done

for file in *sorted.bedgraph; do (python ∼/bin/liftover_clean.py $file ${file%.bedgraph}clean.bedgraph &); done for file in *clean.bedgraph; do (/seq/ucsc/bedGraphToBigWig $file ∼/annotations/hg38_chrom.sizes

${file%sortedclean.bedgraph}.bw &); done

2. Use the script eclip_bigwig2bedgraph.py to extract data for each RNA. This script produces one multibedgraph file with all data and all the extracted bedgraph files for each input bw file.

python ∼/bin/eclip_bigwig2bedgraph.py XIST K562 . hsXIST.bed eCLIP_K562_XIST.multibedgraph &

python ∼/bin/eclip_bigwig2bedgraph.py MALAT1 K562 . hsMALAT1.bed eCLIP_K562_MALAT1.multibedgraph &

python ∼/bin/eclip_bigwig2bedgraph.py NEAT1 K562 . hsNEAT1.bed eCLIP_K562_NEAT1.multibedgraph &

python ∼/bin/eclip_bigwig2bedgraph.py MALAT1 HepG2 . hsMALAT1.bed eCLIP_HepG2_MALAT1.multibedgraph &

python ∼/bin/eclip_bigwig2bedgraph.py NEAT1 HepG2 . hsNEAT1.bed eCLIP_HepG2_NEAT1.multibedgraph &

3. Make a header file for all 121 K562 and 103 HepG2 RBP eCLIP experiments.

ls K562_*_XIST_?.bedgraph | sed ’s/_XIST//g’ | sed ’s/\.bedgraph//g’ | sed ’s/_0/_SMInput/g’ | sed ’s/_1/_eCLIP1/g’ | sed ’s/_2/_eCLIP2/g’ | tr ’\n’ ’\t’ > header_K562_363.txt

ls HepG2*MALAT1_?.bedgraph | sed ’s/_MALAT1//g’ | sed ’s/\.bedgraph//g’ | sed ’s/_0/_SMInput/g’ | sed ’s/_1/_eCLIP1/g’

| sed ’s/_2/_eCLIP2/g’ | tr ’\n’ ’\t’ > header_HepG2_309.txt

4. Then use the eclip_normalize.py to normalize against all controls. Given that the values in the bedgraph files are not the read numbers, I took the average of all the files as the background (121 for K562 and 103 for HepG2).

python eclip_normalize.py XIST 100 header_K562_363.txt eCLIP_K562_hsXIST_100nt.multibedgraph

Note, the bigWig files were made using the following scripts from Yeo lab, and essentially each file is normalized as number of reads per million. Then combine all the binned bedgraph files. https://github.com/YeoLab/gscripts/blob/master/gscripts/general/make_bigwig_files.py https://github.com/YeoLab/gscripts/blob/master/gscripts/general/normalize_bedGraph.py

The output files from steps 2-4 have the following dimensions:

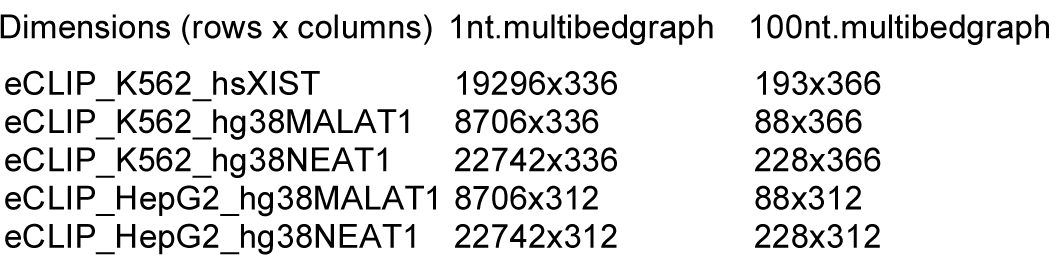

4. Note, editing large pdf or svg files in Illustrator is very slow. Here are some tricks to improve the performance. https://helpx.adobe.com/illustrator/kb/optimize-illustrator-performance-mac-os.html. Make a pdf file in Illustrator to store the names of the 121 samples for K562, and 103 samples for HepG2. This will replace the long file names for the tracks.

tr ’\t’ ’\n’ < header_K562_363.txt | grep eCLIP1 | sed ’s/_eCLIP1//g’

tr ’\t’ ’\n’ < header_HepG2_309.txt | grep eCLIP1 | sed ’s/_eCLIP1//g’

6. The normalized matrix (*normmatrix) files were clustered using the city-block distance and single-linkage, and visualized in TreeView, exported at x:2 y:1.

Reasons for choosing the parameters in clustering are as follows.

A. Although the vectors to be clustered are ratios, no log transformation is used because only the positive enrichment is meaningful.

B. Only arrays (here RBP eCLIP profiles) are normalized, because assessing the overall pattern similarity is high priority in clustering.

C. The arrays are not centered, again because only positive enrichment is meaningful

D. The correlation based similarity metrics are not appropriate because the magnitude of enrichment matters in this calculation.

E. The downside of the Euclidean or city-block distance is that similar patterns may be separated due to the difference in magnitude.

7. Then use the pc2track.py command to make principal component tracks for visualization. The top seven tracks were shown for the three lncRNAs

python pca2tracks.py eCLIP_K562_hsXIST_all121_100nt_pca_array.pc.txt 7 array eCLIP_K562_hsXIST_all121_100nt_pca_array

python pca2tracks.py eCLIP_K562_hg38MALAT1_all121_100nt_pca_array.pc.txt 7 array eCLIP_K562_hg38MALAT1_all121_100nt_pca_array

python pca2tracks.py eCLIP_K562_hg38NEAT1_all121_100nt_pca_array.pc.txt 7 array eCLIP_K562_hg38BEAT1_all121_100nt_pca_array

8. Use this script to convert the PARIS data in bed format to chr1 based mature transcript: genome2transcript.py. This can be used to compare with the eCLIP data.

#### Analysis of RBP motifs based on eCLIP

RBP motifs were derived from previous publications. These motifs are readily available from previous publications and mapped to the human XIST RNA: HNRNPK (CCCC), KHSRP (CCCC), TARDBP (GTRTG), RBM22 (CGG) (Dominguez et al., 2018; Van Nostrand et al., 2017).

#### Analysis of mouse NPC PIRCh data

1. The PIRCh paired end reads were first combined to form gapped reads as follows before mapping. The petwo2gap.py script makes the reverse complement of read 2 in each pair and append that to the end of read 1.

python petwo2gap.py NPC_H3K4Me3_rep1_R1.fastq NPC_H3K4Me3_rep1_R2.fastq NPC_H3K4Me3_rep1.fastq &

2. Map reads to mm10. Many of the mapped reads have insertions (I in the CIGAR string), suggesting that the two reads are overlapping on each fragment. The mapping statistics are included as follows for both NPC and human FL3 PIRCh data.

for file in NPC*.fastq.gz; do (star-static --readFilesCommand gunzip -c --runMode alignReads --genomeLoad LoadAndKeep --outFilterScoreMinOverLread 0.33 --outFilterMatchNminOverLread 0.33 --scoreGapNoncan 0 --scoreGapGCAG 0 --scoreGapATAC 0 --scoreInsOpen 0 --scoreInsBase 0 --alignSplicedMateMapLminOverLmate 0.33 -- genomeDir starmm10 --readFilesIn npc_fastq/$file --outFileNamePrefix npc_fastq/${file%.fastq.gz}_mm10 -- outReadsUnmapped Fastx --outFilterMultimapNmax 1 --outSAMattributes All --alignIntronMin 1 --chimSegmentMin 15 -- chimJunctionOverhangMin 15 --limitOutSJcollapsed 3000000 --runThreadN 8 &); done

| Samples | Total | Unique and (%) | Too many loci and (%) |
| --- | --- | --- | --- |
| NPC_H3K27Ac_rep1.fastq.gz | 49789075 | 38554298 77.44% | 7263690 14.59% |
| NPC_H3K27Ac_rep2.fastq.gz | 74126049 | 56386602 76.07% | 11958468 16.13% |
| NPC_H3K27Me3_rep1.fastq.gz | 58730198 | 40141231 68.35% | 9304105 15.84% |
| NPC_H3K27Me3_rep2.fastq.gz | 50097333 | 38159419 76.17% | 7995706 15.96% |
| NPC_H3K4Me3_rep1.fastq.gz | 47341615 | 36277437 76.63% | 7268422 15.35% |
| NPC_H3K4Me3_rep2.fastq.gz | 69344567 | 53836981 77.64% | 11111068 16.02% |
| NPC_IgG_rep1.fastq.gz | 46986293 | 38158603 81.21% | 6766164 14.40% |
| NPC_IgG_rep2.fastq.gz | 59936890 | 47485939 79.23% | 8639650 14.41% |
| NPC_Input_rep1.fastq.gz | 50582511 | 41505853 82.06% | 7820731 15.46% |
| NPC_Input_rep2.fastq.gz | 62252561 | 51160069 82.18% | 9779852 15.71% |

3. Convert all files to bam.

for file in NPC*sam; do (samtools view -bS -o ${file%out.sam}bam $file &); done

for file in NPC*bam; do (samtools sort -o ${file%.bam}_sorted.bam $file &); done

for file in NPC*sorted.bam; do (samtools index $file &); done

4. Convert Xist mapped reads back to fastq. Note the redirection is different for the two files.

for file in *Aligned_sorted.bam; do (samtools view $file chrX:103460373-103483233 | awk ’{print "@" $1 "\n" $10 "\n+\n" $11}’ > ${file%Aligned_sorted.bam}Xist.fastq &); done

for file in *Chimeric_sorted.bam; do (samtools view $file chrX:103460373-103483233 | awk ’{print "@" $1 "\n" $10 "\n+\n" $11}’ >> ${file%Chimeric_sorted.bam}Xist.fastq &); done

5. Given that the combined paired end reads would have insertions (as defined by the SAM CIGAR tag “I”) due to the partial overlap of the two reads, I edited the fastq files to make each read shorter, e.g. to 40nt, so that the insertions will be avoided. This processing is performed as follows: for file in *mm10Xist.fastq; do (cut -c1-40,113-152 $file > ${file/Xist/Xist80nt} &); done

6. Map Xist reads to mmXist with the following specific parameters. For mapping to the small genome, the scoring system was changed to increase penalty from 0 to −30 for gap opening --scoreGap. for file in NPC*Xist80nt.fastq.gz; do (star-static --readFilesCommand gunzip -c --runMode alignReads --genomeLoad LoadAndKeep –outFilterScoreMinOverLread 0.33 --outFilterMatchNminOverLread 0.33 --scoreGap -30 --scoreGapNoncan 0 --scoreGapGCAG 0 --scoreGapATAC 0 -- scoreInsOpen 0 --scoreInsBase 0 --alignSplicedMateMapLminOverLmate 0.33 --genomeDir starmmXist --readFilesIn npc_fastq/$file --outFileNamePrefix npc_fastq/${file%10Xist80nt.fastq.gz}Xist --outReadsUnmapped Fastx -- outFilterMultimapNmax 1 --outSAMattributes All --alignIntronMin 1 --chimSegmentMin 15 --chimJunctionOverhangMin 15 --limitOutSJcollapsed 3000000 --runThreadN 8 &); done

7. Use samPairingCalling.test.pl to convert the chiastic reads to normal gapped reads (Lu et al., 2016). This step is only used to combine the two files, not to assemble the duplex groups. Instead given that only three major groups are discernable, reads are grouped by their span.

for file in $(ls npc_fastq/*mmXistAligned.out.sam | cut -d’/’ -f7); do (perl samPairingCalling.test.pl -i npc_fastq/$file -j npc_fastq/${file%Aligned.out.sam}Chimeric.out.junction -s npc_fastq/${file%Aligned.out.sam}Chimeric.out.sam -o npc_fastq/${file%Aligned.out.sam}_geometric -g starmmXist/mmXist.fa -z starmmXist/chrNameLength.txt -a empty.gtf -t starmmXist/mmXist.fa -l 15 -p 2 -c geometric 1>npc_fastq/${file%Aligned.out.sam}_geometric.stdout 2>npc_fastq/${file%Aligned.out.sam}_geometric.log &); done

10. Make bedgraph for visualization on IGV. The bedgraph files can be batch loaded from Finder, using the search function. The following is the script for automated processing of all samples.

for file in *geometric.sam; do (samtools view -bS -o ${file%sam}bam $file; samtools sort ${file%sam}bam

${file%.sam}_sorted; samtools index ${file%.sam}_sorted.bam; genomeCoverageBed -ibam ${file%.sam}_sorted.bam -bg

-split -g mmXist.size > ${file%sam}bedgraph &); done

11. To normalize the PIRCh data, use the script pirch_normalize.py. Input: individual bedgraph files directly derived from the bam data, and a header file of all the samples (pirch_NPC_header.txt, e.g. NPC_Input_rep1). Output: normalized in 100nt windows.

python pirch_normalize.py hg38XIST 100

for file in *norm100nt.bedgraph; do (sed ’s/hg38/hs/g’ $file > ${file/mm10/mm} &); done

12. Take the geometric mean for each pair of duplicates to remove the noise. For example

paste NPC_H3K27Ac_rep?_mmXist_geometric_norm100nt.bedgraph | awk ’{print $1, ’\t’, $2, ’\t’, $3, ’\t’, ($4*$8)**0.5}’ > avg_NPC_H3K27Ac_mmXist_geometric_norm100nt.bedgraph

13. Then the PIRCh data are lifted to the hsXIST coordinates to facilitate comparison with the human HNRNPU fRIP, eCLIP, and other related data:

for file in avg*; do (liftOver -minMatch=0.2 -minBlocks=0.2 -fudgeThick $file mmtohsXIST.liftoverchain

${file/.bedgraph/_hsXIST.bedgraph} unmapped &); done

#### Analysis of human FL3 PIRCh data

1. The following three FL3 PIRCh data files were obtained in bigwig format mapped to hg19: FL3_H3_PA_nugen.bw, FL3_IgG_PA_nugen.bw and FL3_Input_nugen.bw. The data were lifted to hg38 using the liftOver utility from UCSC.

2. RNAs were extracted from the bw files using bigWigToBedGraph:

for file in FL3*hg38.bw; do (bigWigToBedGraph -chrom=chrX -start=73820650 -end=73852753 $file

${file%.bw}_XIST.bedgraph &); done

3. Normalization of the FL3 PIRCh data was performed using the same method described above (pirch_normalize.py).

#### PARIS experiments in mES cells

The PARIS protocol was performed as previously described with slight modifications (Lu et al., 2016). HATX mES cells were treated with 0.5mg/ml AMT (Sigma) and crosslinked with 365nm UV for 30min in Stratalinker 2400. Cell lysate was digested with S1 nuclease and RNA purified using TRIzol, and further fragmented with ShortCut RNase III. RNA was separated by 10% native polyacrylamide gel and then the first dimension gel slices were further electrophoresced in a second dimension 20% urea-denatured gel. Crosslinked RNA above the main diagonal was eluted and proximity ligated with T4 RNA Ligase I. After ligation, samples were denatured and purified using Zymo RNA Clean & Concentrator, and photo-reversed with 254 nm UV for 5 min. Proximity-ligated RNA molecules were then ligated to barcoded adapters and converted to sequencing libraries. Libraries were sequenced on the Illumina NextSeq.

#### Analysis of mouse PARIS data

1. Sequencing was performed on a NextSeq together with PHIX spikein. For this PARIS library, PHIX reads were removed based on barcode information. Two sequencing runs were performed for the PARIS libraries, generating hatx1.fastq and hatx2.fastq. Then duplicates were removed: /home/zhipeng/bin/readCollapse -U hatx.fastq -O hatx_nodup.fastq &

2. Trim the 5’ and 3’ end adapter sequence:

java -jar trimmomatic-0.32.jar SE -threads 16 -phred33 hatx_nodup.fastq hatx_trim_nodup.fastq ILLUMINACLIP:P6SolexaRC35.fa:3:20:10 HEADCROP:16 MINLEN:20 &

3. Quality of the processed files was visualized using fastqc.

4. Map reads to the mm10 reference. Given that mouse Xist contains multiple nearly identical repeats, I allowed multiple mapping in this run (--outFilterMultimapNmax 20).

star-static --runMode alignReads --runThreadN 16 --genomeDir starmm10 --genomeLoad LoadAndKeep --readFilesIn hatx_trim_nodup.fastq --limitOutSJcollapsed 3000000 --outFileNamePrefix hatx_trim_nodup_mm10 -- outReadsUnmapped Fastx --outSAMattributes All --outFilterMultimapNmax 20 --outFilterScoreMinOverLread 0.33 -- outFilterMatchNminOverLread 0.33 --scoreGapNoncan 0 --scoreGapGCAG 0 --scoreGapATAC 0 --scoreInsOpen 0 -- scoreInsBase 0 --alignIntronMin 1 --alignSplicedMateMapLminOverLmate 0.33 --chimSegmentMin 15 -- chimJunctionOverhangMin 15

5. Convert Xist mapped reads back to fastq. Note the redirection is different for the two files. Since that some of the reads are output to both the Aligned.out.sam and Chimeric.out.sam files, we need to take the unique reads for mapping to mmXist.

samtools view hatx1_trim_nodup_mm10Aligned_N_sorted.bam chrX:103460373-103483233 | awk ’{print "@" $1 "\n" $10 "\n+\n" $11}’ > hatx_mm10Xist1.fastq

samtools view hatx1_trim_nodup_mm10Chimeric_sorted.bam chrX:103460373-103483233 | awk ’{print "@" $1 "\n" $10 "\n+\n" $11}’ > hatx_mm10Xist2.fastq

samtools view hatx2_trim_nodup_mm10Aligned_sorted.bam chrX:103460373-103483233 | awk ’{print "@" $1 "\n" $10 "\n+\n" $11}’ > hatx_mm10Xist3.fastq

samtools view hatx2_trim_nodup_mm10Chimeric_sorted.bam chrX:103460373-103483233 | awk ’{print "@" $1 "\n" $10 "\n+\n" $11}’ > hatx_mm10Xist4.fastq

cat hatx_mm10Xist* > hatx_mm10Xist.fastq

awk ’NR%4 {printf "%s ",$0;next}1’ hatx_mm10Xist.fastq > a; sort a | uniq -u > b awk ’{print $1 "\n" $2 "\n" $3 "\n" $4}’ b > hatx_mm10Xist_uniq.fastq

6. Map Xist reads to mmXist with the following specific parameters. For mapping to the small mmXist “genome”, the -- scoreGap option was changed from 0 to -30.

star-static --runMode alignReads --genomeLoad LoadAndKeep --outFilterScoreMinOverLread 0.33 -- outFilterMatchNminOverLread 0.33 --scoreGap -10 --scoreGapNoncan 0 --scoreGapGCAG 0 --scoreGapATAC 0 --

scoreInsOpen 0 --scoreInsBase 0 --alignSplicedMateMapLminOverLmate 0.33 --genomeDir starmmXist --readFilesIn hatx_mm10Xist_uniq.fastq --outFileNamePrefix hatx_mmXist_uniqp10 --outReadsUnmapped Fastx -- outFilterMultimapNmax 1 --outSAMattributes All --alignIntronMin 1 --chimSegmentMin 15 --chimJunctionOverhangMin 15 --limitOutSJcollapsed 3000000 --runThreadN 8

7. Use samPairingCalling.test.pl to convert the chiastic reads to normal gapped reads.

perl samPairingCalling.test.pl -i hatx_mmXist_uniqp10Aligned.out.sam -j hatx_mmXist_uniqp10Chimeric.out.junction -s hatx_mmXist_uniqp10Chimeric.out.sam -o hatx_mmXist_uniqp10geometric -g starmmXist/mmXist.fa -z

/home/zhipeng/starmmXist/mmXist.size -a empty.gtf -t starmmXist/mmXist.fa -l 15 -p 2 -c geometric 2>hatx_mmXist_uniqp10geometric_log.txt

8. Convert the output DG information to bed format for visualization in IGV.

samtools view -bS -o hatx_mmXist_uniqp10geometricbam hatx_mmXist_uniqp10geometricsam

samtools sort hatx_mmXist_uniqp10geometricbam hatx_mmXist_uniqp10geometric_sorted

samtools index hatx_mmXist_uniqp10geometric_sorted.bam

python dg2bed.py hatx_mmXist_uniqp10geometric hatx_mmXist_uniqp10geometric.bed bed12

9. To compare with the human XIST structure, I lifted the mouse coordinates to the human one hsXist. Of the 203 DGs (including ones with only identical reads), 108 can be lifted to human XIST coordinates using the adjusted liftOver parameters shown as follows based on previous tests (Lu et al., 2016).

liftOver -minMatch=0.2 -minBlocks=0.2 -fudgeThick hatx_mmXist_uniqp10geometric.bed mmtohsXIST.liftoverchain hatx_mmXist_uniqp10geometric_hsXIST.bed unmapped

10. To compare the mouse lifted PARIS data with the PARIS data from HEK293 cells, use the following scripts. This shuffling test (1000 times) showed that P<0.001.

cp AMT_Stress_trim_nodup_starhsXIST_l15p2_NGmin1386_arc.bed a12.bed

cp hatx_mmXist_uniqp10geometric_hsXIST.bed b12.bed

comparehelix.sh a12.bed b12.bed

dgshuffle.sh hsXISTsimp.bed /hsXIST.size commonlist

11. DG66 and DG69 in the mouse Xist PARIS data are consistent with the human XIST PARIS data (Lu et al., 2016), one of the most conserved long-range duplex. These two DGs were extracted as follows.

awk ’($21∼/DG:i:66/)||($21∼/DG:i:69/)’ hatx_mmXist_uniqp10geometricsam | cut -f4,6,10 > hatx_DG66DG69

#### Generation of knockin cell lines

To test the role of the mouse *Xist* architecture in m6A modification specificity, the A-repeat domain was relocated to several regions in the mouse *Xist* gene using the CRISPR-Cas9 system. Synthesized, HPLC purified sgRNAs were purchased through Synthego. Donor plasmids were generated through overlap extension PCR of 3 fragments: the A-repeat sequence flanked by two 800 bp homology arms to each respective genomic region. The resulting PCR product was then ligated into the pCR-Blunt-II-TOPO vector, using the Zero Blunt TOPO PCR Cloning Kit according to manufacturer’s instructions. Cas9 protein with NLS (PNA Bio) was complexed with sgRNAs in microfuge tubes for 10 min at 37C, and immediately transferred to ice. 1×10^6 TXY:dSX cells were nucleofected with pre-complexed CRISPR RNP and 20 ug of donor plasmid using an Amaxa nucleofector with the manufacturer’s recommended settings for mES cells. Cells were then grown for 72h before low-density splitting into single-cell colonies. Colonies were picked using a light dissection scope and grown in 48-well plates to establish clonal populations. For genotype screening, genomic DNA was extracted using QuickExtract (Lucigen), and then subject to PCR with primers spanning the knockin site (see Supplemental Table for details). Clones were screened by looking for a PCR product significantly higher in size than in that of non-targeted TXY: ΔSX cells. PCR products were then Sanger sequenced through Stanford ELIM using the forward PCR primer as a sequencing primer to verify knockin of the A-repeat.

To make the guide RNAs, DNA template was designed as follows: T7_promoter + sgRNA + scaffold (GAAATTAATACGACTCACTATAGG [sgRNA] GTTTTAGAGCTAGAAATAGCAAGTTAAAATAAGGCTAGTCCGTTATCAACTTGAAAAAGTGGCACCGAGTCGGTGCTTTT). the following single-strand DNA were ordered from IDT DNA and used to make the duplex DNA for in vitro transcription.

mmX.5136f (middle of BCD domain): GAAATTAATACGACTCACTATAGGAGTTAGAAAGATGTGACCTGGTTTTAGAGCTAGAAATAGCAAGTTAAAATAAGGCTAGTCCGTTATCAACTTGAAAAAGTGGCACCGAGTCGGTGCTTTT

mmX.5136r (middle of BCD domain): AAAAGCACCGACTCGGTGCCACTTTTTCAAGTTGATAACGGACTAGCCTTATTTTAACTTGCTATTTCTAGCTCTAAAACCAGGTCACATCTTTCTAACTCCTATAGTGAGTCGTATTAATTTC

mmX.14555f (middle of exon6 domain): GAAATTAATACGACTCACTATAGGTAAAGCCGGGACCTAACTGTGTTTTAGAGCTAGAAATAGCAAGTTAAAATAAGGCTAGTCCGTTATCAACTTGAAAAAGTGGCACCGAGTCGGTGCTTTT

mmX.14555r (middle of exon6 domain): AAAAGCACCGACTCGGTGCCACTTTTTCAAGTTGATAACGGACTAGCCTTATTTTAACTTGCTATTTCTAGCTCTAAAACACAGTTAGGTCCCGGCTTTACCTATAGTGAGTCGTATTAATTTC

mmX.17623f (after exon6 domain): GAAATTAATACGACTCACTATAGGTATGTGATCAAAGCAGATGAGTTTTAGAGCTAGAAATAGCAAGTTAAAATAAGGCTAGTCCGTTATCAACTTGAAAAAGTGGCACCGAGTCGGTGCTTTT

mmX.17623r (after exon6 domain): AAAAGCACCGACTCGGTGCCACTTTTTCAAGTTGATAACGGACTAGCCTTATTTTAACTTGCTATTTCTAGCTCTAAAACTCATCTGCTTTGATCACATACCTATAGTGAGTCGTATTAATTTC

The following PCR primers were used to for cloning and testing (LHA: left homology arm, RHA: right homology arm): 5KI_LHA_F: GAGAAAGCTTGACTTCCAGAGACATAGAATTTCACTTTG

5KI_LHA_R: CCCCGATGGGCAAGAATATATAAACAATGAAGGGCGATAGCACCCATGAC

5KI_repA_F: TGTCATGGGTGCTATCGCCCTTCATTGTTTATATATTCTTGCC

5KI_repA_R: ATCTCCATCAGTTAGAAAGATGTGACCTGACTCACAAAACCATATTTCC

5KI_RHA_F: GGTGGATGGAAATATGGTTTTGTGAGTCAGGTCACATCTTTCTAACTG

5KI_RHA_R: GAGAGAATTCTACAAATAAGTCTTCACCAGATG

14KI_LHA_F: GAGAAAGCTTTGCCCAGGTCACATTATG

14KI_LHA_R: CCCCGATGGGCAAGAATATATAAACAATGAAACAGTTAGGTCCCGGCTTTATAG14KI_repA_F: GTTCTATAAAGCCGGGACCTAACTGTTTCATTGTTTATATATTCTTGCC

14KI_repA_R: AGAAAGTAATCACTGTTCACTGATAAAGCCAACTCACAAAACCATATTTCC

14KI_RHA_F: GGTGGATGGAAATATGGTTTTGTGAGTTGGCTTTATCAGTGAACAG

14KI_RHA_R: GAGAGAATTCTATATAATTCTTTAAAAATATTATTCACTCAG

17KI_LHA_F: GAGAAAGCTTTCCTTACTATAATATACTCAAGGTGG

17KI_LHA_R: CCCCGATGGGCAAGAATATATAAACAATGAATGGTAGGATGTGCTTAATTG

17KI_repA_F: ATATTGCTACCAATTAAGCACATCCTACCATTCATTGTTTATATATTCTTGCC

17KI_repA_R: GTACACAGTTCATTTATGTGATCAAAGCAGATGAACTCACAAAACCATATTTCC

17KI_RHA_F: GGTGGATGGAAATATGGTTTTGTGAGTTCATCTGCTTTGATCACATAA

17KI_RHA_R: GAGAGAATTCAGGGCCACTGAGTTAGAAAC

### Genotyping Primers

5KI_Genotype_F and 5KI_Genotype_R: CCAGCCCTGTGTGCATTTAG, AGCCTTATCCAGTGTCCAGG

14KI_Genotype_F and 14KI_Genotype_R: TTCCACCTCCTCAGTCAAGC, TGCTTTGGTGAGGCTCAGTA

17KI_Genotype_F and 17KI_Genotype_R: AGCAAGCCTGACCCTAAAGT, TGGTGGGAAGATGACTCCAG

### sgRNA Test Primers

5KI_gRNA_F and 5KI_gRNA_R: CCAGCCCTGTGTGCATTTAG, GGTTTGATTCCCCAGCACAG

14KI_gRNA_F and 14KI_gRNA_R: GCCTGGTGTGCAATGACTTT, TGGGATTATCTCACTCTGGCC

17KI_gRNA_F, and 17KI_gRNA_R: GAGCCAGGTTGAAGAGGTCT, AACTACCCCACCCACTCAAC

#### MeRIP-Seq in mouse ES cells

Total mES RNA was subjected to one round of Poly(A)Purist MAG treatment to enrich for polyadenylated RNAs as per manufacturer’s instructions (Ambion). RNA was then fragmented to 100nt median sized fragments using RNA Fragmentation Reagents (Ambion) and subjected to one round of m6A immunoprecipitation. For immunoprecipitation of RNA, 5 ug of m6A antibody (Millipore) was coupled to 40 ul Protein A Dynabeads (Novex) in 100 ul 1X IPP Buffer (50mM Tris-HCl, pH 7.5; 150mM NaCl; 0.1% NP-40; 5mM EDTA) overnight at 4C. Beads were then washed twice in 1X IPP Buffer. Fragmented RNA was denatured at 70C for 2 min, cooled on ice, and bound to antibody-beads in 185 ul 1X IPP Buffer for 3h at 4C. Beads were then washed sequentially with (2x) 500 ul 1X IPP Buffer, (2x) 500 ul Low Salt Buffer (0.25X SSPE; 1mM EDTA; 0.05% Tween-20; 37.5mM NaCl), (2x) 500 ul High Salt Buffer (0.25X SSPE; 1mM EDTA; 0.05% Tween-20; 137.5mM NaCl), (1x) 500 ul TET Buffer (10mM Tris-HCl, pH 8.0; 1mM EDTA; 0.05% Tween-20). Beads were eluted with 50 ul RLT Buffer (Qiagen RNeasy Mini Kit) and incubated at 25C for 5 min, and recovered with RNeasy Mini Kit followed by concentrating with Zymo RNA Clean & Concentrator in 10 ul water. 10 ng of input RNA (before immunoprecipitation) and 10 ng of immunoprecipitated RNA were then used to prepare sequencing libraries using SMARTer Stranded RNA-Seq Kit - Pico Input Mammalian (Clontech #634411), as per manufacturer’s instructions. Sequencing libraries were pooled and sequenced on the Illumina MiSeq (files named *mar08* and *apr26*) and NextSeq (files named *jun24*).

#### Global analysis of meRIP-seq data

The following input and immunoprecipitation paired-end (75nt x2 or 76nt x2) datasets were generated using the method described above. Five alleles were analyzed using MiSeq: wt, dsx (ΔSX, or A-repeat deletion), ki5, ki14 and ki17_1. All 6 alleles were analyzed by NextSeq: wt, dsx (ΔSX, or A-repeat deletion), ki5, ki14 and ki17_1 and ki17_2. The first 5 of the NextSeq libraries were re-sequencing of the same libraries for MiSeq. The general pipeline is as follows: map paired-end reads to mm10 convert reads to bam format and separate the two strands convert to bedgraph and bigwig for visualization. Fo r global analysis: use m6Aviewer (Antanaviciute et al., 2017). For specific analysis of Xist: extract data in the mouse Xist region normalize against background ranges normalize against wildtype input in 200nt windows count coverage in each predefined m6A domains plot m6A modification levels in bar graphs compare known m6 A motifs with actual modification sites as determined by meRIP-seq. Detailed analysis pipeline is described as follows.

Convert paired end reads to gap reads map to hg38 extract reads mapped to XIST map to hsXIST ‘minigenome’ assemble mapped Aligned and Chimeric reads ex tract long pairs make distance distribution, bedgraph and arcs f or both short and long pairs.

1. Map paired-end reads to mm10 using the STAR (here *1.fastq indicate the first mate of each pair of paired-end files). for file in *1.fastq; do (star-static --runMode alignReads --genomeLoad LoadAndKeep --genomeDir starmm10/ -- readFilesIn $file ${file/1.fastq/2.fastq} --outFileNamePrefix ${file/1.fastq/_mm10} --outReadsUnmapped Fastx -- outFilterMultimapNmax 1 --outSAMattributes All --alignIntronMin 20 --runThreadN 6 &); done

2. Convert to bam files and separate the strands

for file in *sam; do (samtools view -bS -o ${file/out.sam/bam} $file &); done for file in *bam; do (samtools sort -o ${file/.bam/_sorted.bam} $file &); done for file in *sorted.bam; do (samtools index $file &); done

for file in *_mm10Aligned_sorted.bam; do (

samtools view -b -f 128 -F 16 $file > ${file}fwd1.bam; samtools view -b -f 80 $file > ${file}fwd2.bam;

samtools view -b -f 144 $file > ${file}rev1.bam; samtools view -b -f 64 -F 16 $file > ${file}rev2.bam;

samtools merge ${file/sorted.bam/pos.bam} ${file}fwd1.bam ${file}fwd2.bam;

samtools merge ${file/sorted.bam/neg.bam} ${file}rev1.bam ${file}rev2.bam;

samtools index ${file/sorted.bam/pos.bam}; samtools index ${file/sorted.bam/neg.bam}); done &

3. Convert bam to bedgraph and then bw for visualization. for file in *pos.bam *neg.bam; do (bedtools genomecov -bg - split -ibam $file -g starmm10/chrNameLength.txt > ${file/bam/bedgraph} &); done

for file in *bedgraph; do (bedGraphToBigWig $file starmm10/chrNameLength.txt ${file/bedgraph/bw} &); done

4. Use the m6aViewer software (version 1.6.1) to calculate global changes in m6A modification using the default parameters and negative strand bam files (half the data). To retrieve the columns that contain the dsx or wildtype comparisons, for example, use the following script: head -n 1 All6_differential.txt | awk ’BEGIN {FS="\t"}; { for (i=1; i<=NF;

++i) { if ($i ∼ "dsx") print i } }’

5. Plot the differences using scatter_volcano.py script. Log scale fold changes (LFC) and log scale p values were used as the x-axis and y-axis, respectively. For example: python scatter_volcano.py All6_differential.txt 29 30 m6A_global_scatter_dsx_vs_ki14.png

#### Targeted analysis of Xist m6A modification

1. Continuing from step 3 of the global analysis of meRIP-seq, extract data from the mm10 reference to mmXist coordinates using eclip_bigwig2bedgraph.py and normalize data against input using pirch_normalize.py. python

∼/bin/eclip_bigwig2bedgraph.py mm10Xist bw . 1 mmXist.bed TXYmerip_mmXist.multibedgraph

2. Convert the “chr1” to “mmXist”: awk ’{print "mmXist\t" $2 "\t" $3 "\t" $4}’ XXX_neg_mm10Xist_1.bedgraph > XXX_neg_mmXist_1.bedgraph

3. For normalization, use the following ranges as background: 2000-4500, 5500-9000, 13000-14000 and 16000-17000. The files are normalized so that the wildtype input is set as the baseline. for file in *all*mm10Xist*bedgraph; do (echo $file; awk ’($2>=2000)&&($2<4500)||($2>=5500)&&($2<9000)||($2>=13000)&&($2<14000)||($2>=16000)&&($2<17000) {sum+=$4}; END {print sum}’ $file); done

4. Normalize each file so that coverage in the background ranges are the same as the wildtype input: awk ’{print "mmXist\t" $2 "\t" $3 "\t" $4*norm_factor}’ XXX_inall_mm10Aligned_neg_mm10Xist_1.bedgraph > XXX_inall_mm10Aligned_neg_mmXist_normbgrd.bedgraph

5. To visualize the bedgraph files in IGV, means were taken in 50 nt windows to minimize file size.

6. Count the reads in each predefined domain for making the bar graph of m6A levels. The domains were defined as follows: m6AD1: 0-1400, m6AKI5: 4500-5500, m6AD2: 9100-9900, m6AD3:11400-12300, m6AKI14:14100-15300, m6AD4/m6AKI17: 17000-17900. for file in *mmXist_normbgrd.bedgraph; do (bedtools map -c 4 -o sum -null 0 -a mmXist_m6AD.bed -b $file); done > mmXist_m6AD.count

7. m6A levels were normalized against wildtype and then plotted in MS Excel.

8. Used the script motif.py to make a track for all the m6A motifs in mmXist and a track of motif density in 300nt windows and 50nt steps. There are 333 DRACH motifs in total in the 17918nt mouse Xist transcript and 360 motifs in the 19296nt human XIST.

python motif.py DRACH mmXist.fa mmXist_DRACH.bed 300 50 mmXist_DRACH_density.bedgraph

#### Analysis of the silencing functions of Xist alleles using the meRIP-seq input data

1. Summarize expression levels of all mouse genes. for file in *inall_mm10Aligned*.bam; do (bedtools coverage -s -split - abam $file -b ∼/annotations/mm10refGenePU.bed > ${file/bam/count} &); done

2. Extract strand-specific count for all the input files and then combine them: for file in *all*pos.count; do (awk ’$6=="+"’ $file > ${file}pos); done

for file in *all*neg.count; do (awk ’$6=="+"’ $file > ${file}neg); done

file in *all*countpos; do (cat $file ${file/pos.countpos/neg.countneg} > ${file%_pos.countpos}.countstrand); done

3. Summarize expression by chromosome. Combine the counts from all samples. paste *inall_mm10Aligned.countstrand | cut -f1-4,13,29,45,61,77,93 > Inputall_mm10Aligned.count

3. Take the ratios for each gene against wt, for genes with counts >=100. awk ’($5>=100)&&($6>=100)&&($7>=100)&&($8>=100)&&($9>=100)&&($10>=100) {print $1 "\t" $2 "\t" $3 "\t" $4 "\t" $5/$10 "\t" $6/$10 "\t" $7/$10 "\t" $8/$10 "\t" $9/$10}’ Inputall_mm10Aligned.count > Inputall_mm10Aligned_xci_violin.ratios

4. Plot the expression difference among all the samples using this script: violin_xci.py. The medians of ratios of expression levels all autosomal genes between mutant and wildtype were set to one for each mutant cell line, and then the medians of ratios for X chromosome genes were calculated. Differences between autosome and X chromosome ratios were assessed using Mann-Whitney U test.

#### Analysis of miCLIP data from Linder et al. 2015 Nature Methods

The miCLIP bedgraph files in GSE63753 were downloaded from GEO and lifted to hg38 using the liftOver tool from UCSC genome brower (Linder et al., 2015). Then the data were lifted to the mature XIST transcript coordinates (without introns) using custom python scripts.

#### irCLIP analysis of SPEN and LBR

The irCLIP experiments were performed as described previously (Zarnegar et al., 2016). Briefly, mES cells with engineered Xist mutations were cultured with standard conditions and induced to express Xist (see earlier description on cell culture). Afterward, cells were lysed, immunoprecipitated with antibodies for these RBPs and treated with S1 nuclease. RNP complexes were resolved on denatured polyacrylamide gels and regions above the protein size were excised for RNA extraction and library preparation.

Sequencing output reads were processed by bbmap to remove duplication on fastq level. Remained reads were trimmed off the 3’ solexa adapter and against sequencing quality q20 by cutadapt (version 2.4). Trimmed reads were mapped first to RNA biotypes with high repetitiveness by bowtie2 (version 2.2.9) to our custom built indexes: rRNAs (rRNAs downloaded from Ensembl GRCm38.p6/mm10 and a full, non-repeat masked mouse rDNA repeat from GenBank accession No.BK000964), snRNAs (from Ensembl GRCm38.p6/mm10), miscRNAs (from Ensembl GRCm38.p6/mm10), tRNAs (from UCSC table browser GRCm38.p6/mm10), RetroGenes V6 (from UCSC table browser GRCm38.p6/mm10) and RepeatMasker (from UCSC table browser GRCm38.p6/mm10). Remained reads were mapped to mouse genome GRCm38/mm10 by STAR (version 2.7.1a) with junction file generated from mRNAs and lncRNAs by Genocode GRCm38.p6/mm10 GTF file. Only reads uniquely mapped to the mouse genome were included in the down-stream analysis. The RBP binding loci as suggested by the irCLIP method, was defined as 1-nt shift to the 5’ end of each mapped read. Each locus was extended 5nts up and downstream to shape a local interval, only intervals overlapped between two replicates were included. Then 5nts were trimmed from each side of the overlapped interval to shape the final cluster. Cluster annotation was processed against the Genocode GRCm38.p6/mm10 GTF file. Reads annotated to Xist gene were re-mapped to the Xist mini-genome. Normalization on Xist was processed in the same way as for m6A data.

### QUANTIFICATION AND STATISTICAL ANALYSIS

In relevant figures, figure legends denote the statistical details of experiments including statistical tests used, kind of replicates and the value of n. Asterisks define degree of significance as described in the figure legends. All Student’s t test and Mann-Whitney U-test were analyzed in two-sided. All the sequencing data were aligned to mouse and human genomes (mm10 and hg38) or custom made mini-genomes (mmXist and hsXIST). Statistical analyses and graphics were performed using Python, R and Microsoft Excel.

### DATA AND SOFTWARE AVAILABILITY

All software used in this study is listed in the Key Recourses Table. Custom scripts for analysis were published in GitHub: https://github.com/zhipenglu/xist_structure. Conservation plot is imported from “100 vertebrates Basewise Conservation by PhyloP” at UCSC genome browser. Normalized bedgraph files for all 121 proteins in eCLIP and 25 proteins in fRIP-seq are available in https://www.dropbox.com/sh/24kbqwxafhhzrli/AAADwDA6gdDOY-hWFoS4-V3aa?dl=0. The custom IGV genome for human XIST mature transcript is available in the same folder as well. All raw sequencing reads and raw count matrices generated in this study are available through Gene Expression Omnibus (GEO) with accession number GSE126715 (m^6^A RIP-seq and irCLIP on A-repeat relocation alleles), GSE126716 (PARIS in mouse ES cells).

